# Genotype-associated transcriptional differences among grapevine lines carrying single and pyramided *Rpv* resistance loci

**DOI:** 10.1101/2025.11.27.690962

**Authors:** Martin Hádlík, Miroslav Baránek, Kateřina Baránková, Viera Kovacova

## Abstract

**Background:** Downy mildew, caused by *Plasmopara viticola*, is a major threat to grapevine production. Pyramiding resistance loci is a strategy for improving resistance, but how transcriptional responses differ among genotypes carrying different locus combinations remains unclear. We generated 36 time-resolved transcriptomes from grapevine genotypes carrying *Rpv12*, *Rpv12+1*, or *Rpv12+1+3* and a susceptible control, analyzed differential expression and co-transcriptional networks, and screened protein pairs with AlphaFold2-Multimer.

**Results:** All three resistant genotypes already differed from the susceptible control at the time of inoculation (0 hpi). Genotype differences in the number of differentially expressed genes were largest at 6 hpi, when *Rpv12+1+3* had the most, whereas *Rpv12+1+3* changed least relative to its own 0 hpi state. At 0 hpi, the *Rpv12* genotype examined here had higher transcript levels than the susceptible control for receptor-related genes, including SOBIR1 and homologs of *Arabidopsis* LRIP1, and for EDS1–SAG101–PAD4 family members; neither multilocus genotype showed this pattern, and in direct comparisons most of these genes were higher in *Rpv12* than in *Rpv12+1+3*. A structural screen based on the PN40024 v4 gene models predicted ten receptor-associated interactions, but none was reproduced with longer gene models of the same loci from newer annotations, which differed substantially from the v4 models. Predicted EDS1–SAG101 and EDS1–PAD4 complexes with complete gene models were consistent with known complexes.

In an independent *Rpv12*-carrying transcriptome, about three-quarters of the genes responding in both studies changed in the same direction at early and late time points, but these shared genes reflected general stress- and growth-related functions, and the comparison does not establish *Rpv12* specificity.

**Conclusions:** The examined breeding genotypes differ in their transcriptional state at inoculation and in their temporal response, so genotype, and not only the locus combination, should be treated as a primary variable in transcriptomic studies of grapevine resistance. Structural interaction predictions for leucine-rich repeat receptors depend on the gene models used and require verified gene models before they can guide experiments.

## BACKGROUND

The oomycete *Plasmopara viticola* poses a significant threat to grapevine (*Vitis* spp.), causing devastating downy mildew that can destroy up to 75% of the crop in a single season [1, 2]. While copper-based fungicides remain effective, their intensive use raises environmental concerns, making the cultivation of resistant interspecific genotypes a more ecologically sound approach [1, 3]. Numerous loci of resistance to *Plasmopara viticola* (*Rpv*) have been described, including *Rpv36* and *Rpv37* reported in 2024 [4], with *Rpv1*, *Rpv3*, and *Rpv12* considered major loci most commonly used in breeding programs [5]. Pyramiding resistance loci is a central strategy for durable field resistance [6–8], yet how stacked resistance loci influence early immune signaling remains poorly understood. The *Rpv1* locus (from *Muscadinia rotundifolia*) is considered dominant [9]. *Rpv3* (from *Vitis rupestris*) is a major dominant locus that triggers a hypersensitive response and reduces, but does not halt, pathogen growth [10], and *Rpv12* (from *Vitis amurensis*) is a major dominant locus associated with a hypersensitive response [11, 12]. These *Rpv* loci contain NLR (nucleotide-binding leucine-rich repeat) genes that differ in domain architecture: TIR-NB-LRR (TNL) genes have been described at *Rpv1* and *Rpv3*, whereas the *Rpv12* locus contains a cluster of CC-NB-LRR (CNL) candidate genes [9, 12–14]; these loci were characterized in other accessions. NLR proteins are broadly classified by N-terminal domain into TNLs, CNLs and RPW8-NLRs (RNLs), each followed by NB-ARC and LRR domains [15]. In *Arabidopsis*, TNLs require EDS1 for signaling, whereas many CNLs signal largely independently of EDS1, although EDS1–PAD4 dimers with ADR1 helper NLRs promote a basal immunity branch through which both TNL and CNL receptors signal [16–18]. However, the specific signaling architecture of *Vitis* NLRs and their coordination with cell-surface pattern recognition receptors (PRRs) have not been systematically characterized.

Plant immunity involves two interconnected tiers: pattern-triggered immunity (PTI), activated when cell-surface PRRs detect conserved pathogen-associated molecular patterns (PAMPs), and effector-triggered immunity (ETI), activated when intracellular NLRs detect pathogen effectors [19]. Despite activation in distinct cellular compartments, PTI and ETI overlap in various steps and converge in downstream outputs [20–22]; for example, the signaling hub EDS1 can activate both PTI and ETI outputs and forms EDS1–PAD4 and EDS1–SAG101 heterodimers whose contributions to immune outputs overlap and depend on the receptor and pathogen context [17, 23]. At the cell surface, receptor-like proteins (RLPs) lacking a kinase domain signal through the adaptor kinase SOBIR1 and the co-receptor BAK1 [24, 25]. Grapevine pattern-recognition receptors have been characterized in the LysM-RLK and LRR-RLK families [26, 27], but the RLP–SOBIR1 axis has not been examined.

*Rpv1* and *Rpv3* are associated with transcriptional signatures including calcium signaling, WRKY and MYB activation, ROS production, stilbene biosynthesis, and programmed cell death (PCD) [3, 13, 28], while *Rpv12*, whose locus contains CNL candidate genes, shows earlier PCD onset and metabolite alterations that mirror but often exceed *Rpv3* responses [29, 30]. A *P. viticola* isolate characterized by Gouveia et al. overcame *Rpv3.1*- and *Rpv3.2*-mediated resistance but not *Rpv12*-mediated resistance in the material tested [31]. In addition, recent evidence indicates that the genetic background in which an *Rpv* locus is deployed is a determinant of the resistance outcome rather than a nuisance factor. The same pyramided loci confer beneficial, neutral or detrimental quantitative effects depending on the combination, the background and the pathogen strain [32]; varieties carrying identical *Rpv1*–*Rpv3.1* haplotypes differ in hypersensitive-response frequency, which may reflect their pedigree and introgression history [5]; the intensity of *Rpv3*-dependent resistance varies substantially among carriers of the same haplotype [33]; and a specific *Rpv3-3* haplotype confers resistance through stilbenoid induction rather than the hypersensitive-response-based mechanisms typical of other *Rpv* loci [34].

Breeding programs increasingly deploy pyramided *Rpv* combinations [5, 32], yet whether combinations such as *Rpv12+1* or *Rpv12+1+3* alter immune timing, coordination, and regulatory architecture during the earliest detectable stages of pathogen exposure has not been systematically characterized. The selection process for a new grape variety takes about 15 years [35], and resistance loci are introgressed through multiple generations of backcrossing; consequently, the resulting breeding lines are not strictly isogenic and may carry additional linked genomic regions beyond the targeted loci. This complicates causal attribution of phenotypes to individual loci. Here, we use time-resolved RNA sequencing to characterize transcriptomic responses in genotypes carrying single (*Rpv12*), double (*Rpv12+1*), or triple (*Rpv12+1+3*) resistance loci, compared to a susceptible control, and use structural prediction to examine possible interactions among proteins encoded by genes that differ between genotypes. In the *Rpv12*-carrying genotype, transcripts of LRIP1/SOBIR1-related receptor genes and of EDS1–SAG101 paralogs are elevated at inoculation; the predicted receptor interactions, however, depended on the gene models used and were not reproduced with models from newer annotations. Multilocus genotypes (*Rpv12+1*, *Rpv12+1+3*) show transcriptional patterns that differ from those of *Rpv12* alone, though functional interpretation requires quantification of phenotypic resistance.

## RESULTS

### Experimental design and RNA sequencing overview

To investigate how early transcriptional patterns differ among genotypes carrying different resistance-locus combinations, we performed a controlled infection experiment followed by time-resolved RNA sequencing of leaf tissue. Three grapevine genotypes carrying single (*Rpv12*), double (*Rpv12+1*), or triple (*Rpv12+1+3*) resistance loci, together with a susceptible control, were inoculated with *Plasmopara viticola*. Leaf discs were sampled at 0, 6, and 24 h post-inoculation (hpi), yielding 36 RNA-seq libraries (3 replicates × 4 genotypes × 3 time points). Sampling focused on key transition phases (recognition, early response, and later transcriptional phases) previously defined in grapevine–*Plasmopara* interactions [29, 30]. The experimental design and analytical workflow are summarized in Figure 1. After adapter trimming and quality filtering, all 36 samples passed quality thresholds, and 26,169 protein-coding genes with at least 15 reads in total were retained.

**Figure 1:**
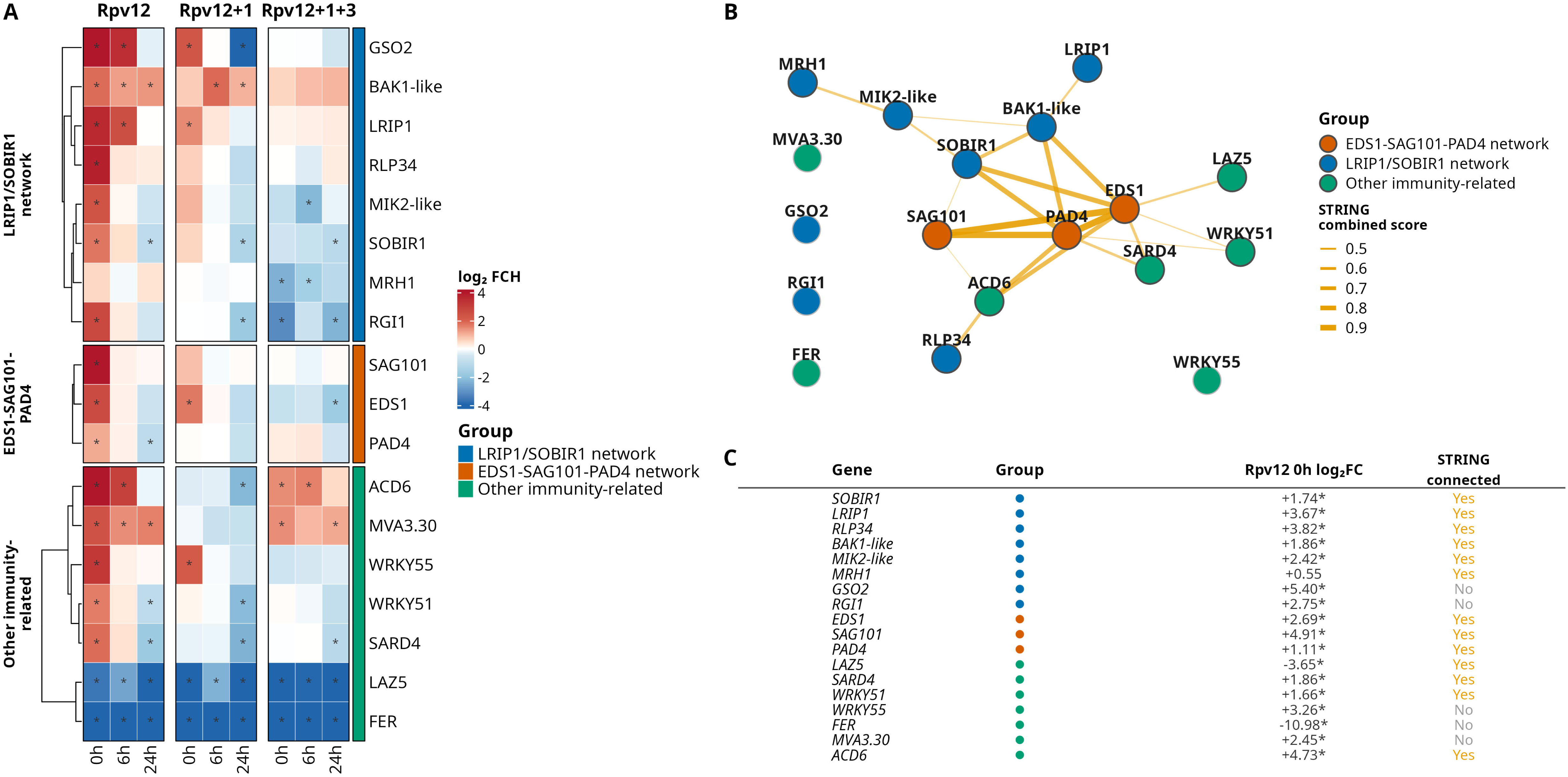
Experimental workflow for artificial inoculation and transcriptomic analysis of four grapevine genotypes. Color code: *Rpv12* – goldenrod; *Rpv12+1* – salmon; *Rpv12+1+3* – cornflower blue; and Pinot Noir – dim gray; as a susceptible control. For each genotype, three independent plants were used as biological replicates. At each time point (0, 6, and 24 hpi), one disc (white circle) was excised from each of three inoculated leaves per plant; leaves from the same plant were processed together to yield one RNA-seq library. In total, 36 libraries were sequenced (4 genotypes × 3 time points × 3 biological replicates). Artificial inoculation was performed by applying a 20 µl drop of spore suspension to the abaxial side of each leaf. Collected samples underwent total RNA isolation, RNA concentration and integrity (RIN) assessment, followed by RNA sequencing and subsequent bioinformatic evaluation.

Sequencing was performed across two batches. Three of four read quality metrics (Additional file 1: Table S1) differed significantly between batches — forward-only survival (FDR = 0.003), read pairs with both mates retained (FDR = 0.004), and reverse-only survival (FDR = 0.014) — reflecting higher adapter trimming rates in one batch (Additional file 2: Figure S1), consistent with known technical variation in multi-run RNA-seq datasets [36]. However, both mates were retained for 88–97% of read pairs in every library, and the proportion of dropped reads did not differ between batches (FDR = 0.077), indicating similar overall data quality. Trimmed reads were aligned to the *Vitis vinifera* PN40024.v4 reference genome [37], yielding median uniquely mapped-read proportions of 90.4-94.3% across genotypes. In a quasibinomial GLM of the uniquely mapped-read proportion with sequencing batch, time point and genotype as predictors, genotype was not associated with mapping efficiency (F = 2.10, p = 0.12), whereas batch (p < 0.001) and time point (p < 0.001) were.

Uniquely mapped-read proportions in this study (Additional file 1: Table S1; mean 91.5%, range 83.5-95.3%) were comparable to two independent grapevine RNA-seq datasets reprocessed for cross-study comparison (Chitarrini et al. [29]: mean 88.4%, range 75.0-92.1%; Shi et al. [38]: mean 93.1%, range 89.4-96.4%), indicating that mapping efficiency in this study was not unusual relative to independently generated grapevine transcriptomes.

To limit the influence of the technical structure on biological comparisons, we evaluated seven strategies spanning raw counts, size-factor (median-of-ratios) normalization, and rlog (variance-stabilizing) transformation, each with and without ComBat batch correction (in both condition-protected and unprotected forms), assessing performance by Spearman correlation (ρ) between PC1 scores and batch assignment (Additional file 3: Figure S2). Among the strategies examined, size-factor normalization [39] combined with condition-protected ComBat correction [40, 41] gave the weakest association between PC1 and sequencing batch for amplitude-based analyses (ρ = −0.035, compared with ρ = −0.640 for raw counts), and rlog transformation [39] followed by condition-protected ComBat gave the weakest association for correlation-based analyses, including clustering and network inference (ρ = −0.062). The two corrected matrices also differed in their mean–variance relationship: after rlog transformation, gene-wise variance was essentially independent of mean expression (Spearman ρ = −0.04), whereas the size-factor-normalized values retained a strong dependence (ρ = −0.68). We therefore used rlog values for PCA and correlation-based analyses, and size-factor-normalized values, which keep the original expression amplitudes, for aggregated expression divergence. Genotypes remained separated after correction with both strategies (Additional file 3: Figure S2). These quality-controlled, batch-corrected datasets formed the basis for all downstream analyses.

Genome-wide between-replicate variance, which includes both biological and technical variation, did not increase with major resistance locus dosage (Additional file 4: Figure S3B; overall linear model: F = 0.72, p = 0.63; dosage terms: F = 0.99, p = 0.46; permutation test of the dosage slope: p = 0.76), and mapping efficiency was not lower in the introgressed genotypes (median 92.1–94.3% vs 90.4% in the susceptible control).

Thus, the main technical structure in the data was the sequencing batch, which was corrected or included in all models, and data quality did not differ systematically between genotypes.

### Genome-wide transcriptional differences between genotypes

To summarize genome-wide transcriptional differences between each resistant genotype and the susceptible control, we calculated the aggregated expression divergence (AED; see Methods) [42], the mean squared difference in condition-protected, batch-corrected, size-factor-normalized expression across all genes (Figure 2B, Additional file 5: Table S2). AED is a distance and does not indicate which genes or pathways differ; it is reported without significance tests, because with three libraries per group, the same plants sampled at all time points and ComBat fitted once with the observed condition labels as protected covariates, permutation p-values would be neither informative nor calibrated. At each time point, the distance between each resistant genotype and the susceptible control was AED = 1.21–2.21 (Figure 2B). The differences at 0 hpi were present at the time of inoculation and may reflect genetic-background variation among the non-isogenic breeding lines as well as, or instead of, effects of the resistance loci.

**Figure 2:**
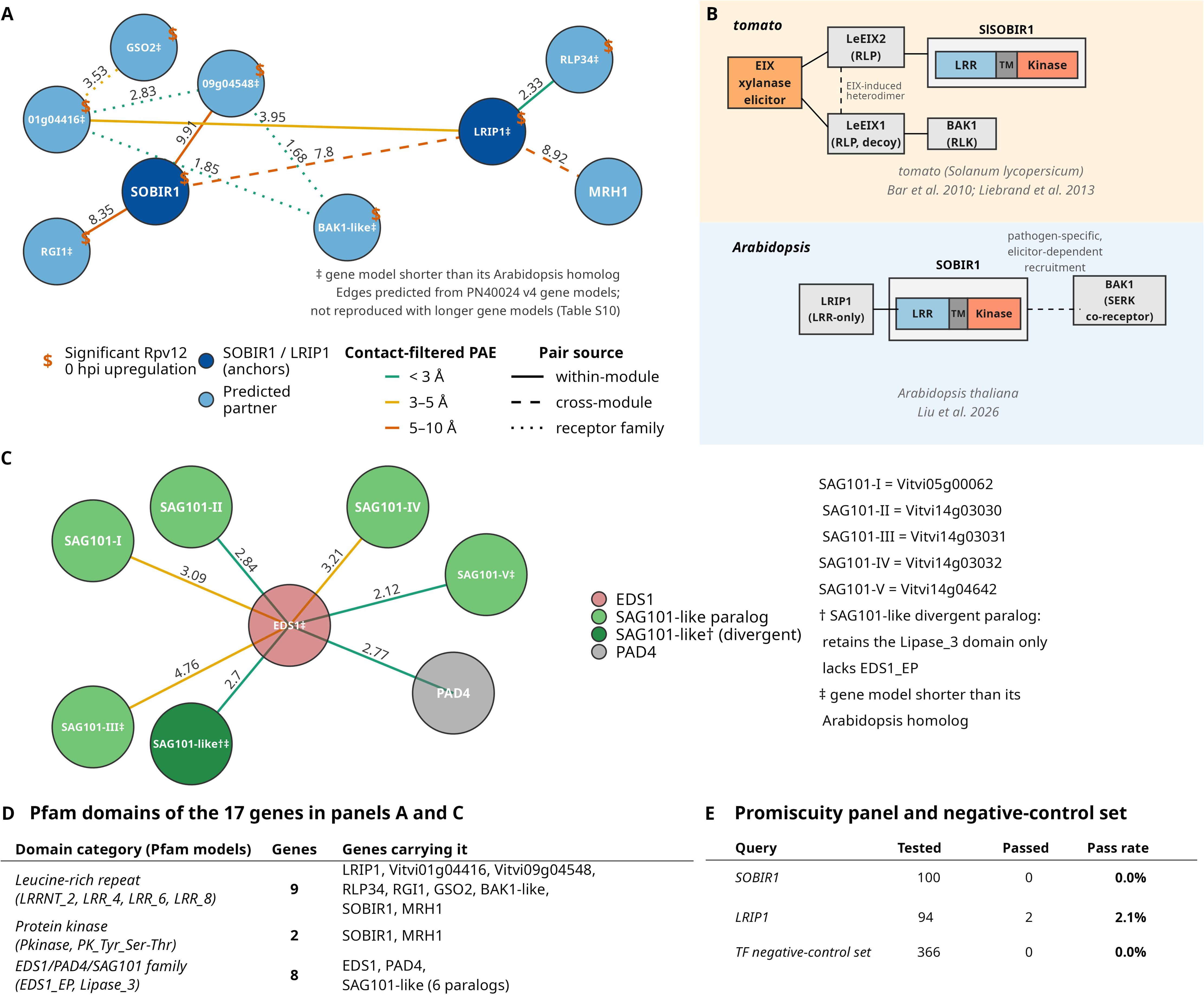
Aggregated expression divergence and principal component structure across single-and multilocus resistance genotypes. (A) Aggregated expression divergence (AED) of each genotype relative to its own 0 hpi state at 6 and 24 hpi (0 hpi = 0 by definition). Points and lines show genotype means (gray, Susceptible; goldenrod, *Rpv12*; salmon, *Rpv12+1*; cornflower blue, *Rpv12+1+3*). Reference magnitudes from external datasets (Additional file 5: Table S2): gray dashed line, two sequencing experiments of an isogenic *Arabidopsis thaliana* line (AED = 0.84; Froussios et al. [43]); green diamonds, mock-treated samples of an independent *Rpv12*-carrying dataset at 12 and 24 h relative to 0 h (0.86 and 0.63; Chitarrini et al. [29]); green dashed line, their mean (0.74). These are reference magnitudes, not significance thresholds. *Rpv12* and *Rpv12+1* lie above them at 6 and 24 hpi; *Rpv12+1+3* and Susceptible lie below. (B) AED of each resistant genotype relative to Susceptible at the same time point (colors and gray dashed line as in A). AED is shown as a descriptive distance; no significance tests were applied (see Methods). (C) PCA (PC1 vs. PC2) of 36 transcriptomes (3 biological replicates × 4 genotypes × 3 time points). Color, genotype (as in A); shape, time point (circle, 0 hpi; triangle, 6 hpi; square, 24 hpi). Axis annotations give selected STRING enrichment terms (UniProt keywords, InterPro domains, STRING local-network clusters) among the top 50 contributing genes per PC (contribution-based, direction-agnostic); they are annotation enrichments, not measurements of activity. PC1 (40.3% variance; signal peptide, UniProt keyword ’Signal’, FDR = 3.35 × 10⁻⁷) separates *Rpv12+1+3* from the other genotypes; PC2 (12.2%; NB-ARC/APAF-1 helical domain annotations typical of NLR proteins, FDR = 6.95 × 10⁻⁶) separates Susceptible from the rest. (D) PCA (PC3 vs. PC4), coding and annotation as in C. PC3 (11.4%; UniProt keyword ’Hypersensitive response’, three genes, FDR = 0.0306) separates *Rpv12+1* from the other genotypes; PC4 (8.17%; mixed STRING local-network cluster including systemic-acquired-resistance (SAR) annotations, three genes, FDR = 0.029) mainly separates Susceptible from *Rpv12* at 24 hpi.

Within genotypes, AED between time points (Figure 2A, Additional file 5: Table S2) was small in *Rpv12+1+3* (0.32–0.54) and in the susceptible control (0.35–0.90), and larger in *Rpv12* (1.20–1.83) and *Rpv12+1* (0.35–1.52). *Rpv12* moved away from its 0 hpi state by 6 hpi and remained at a similar distance at 24 hpi (1.83 and 1.77), whereas *Rpv12+1* moved away by 6 hpi (1.52) and partly back by 24 hpi (1.00).

The genes distinguishing each resistant genotype from the susceptible control also differed in temporal consistency (Additional file 5: Table S2). Of the genes differentially expressed relative to the susceptible control at 0 hpi, 55.7% remained differentially expressed at 24 hpi in *Rpv12+1+3* (Jaccard overlap 0.35–0.49 across time-point pairs), compared with 32.0% in *Rpv12+1* (Jaccard 0.19–0.26) and 28.2% in *Rpv12* (Jaccard 0.14–0.21). Together, *Rpv12+1+3* changed little relative to its own 0 hpi state, whereas *Rpv12* and *Rpv12+1* changed more over time.

To place these magnitudes in context, AED was computed in the same way for external datasets (Additional file 5: Table S2). In an isogenic *Arabidopsis thaliana* line, divisions of the replicates within one experiment gave AED = 0.12–0.22, and two separate sequencing experiments of the same line gave AED = 0.84 [43] (Additional file 6: Table S3). In an independent *Rpv12*-carrying grapevine dataset [29], mock-treated samples differed from their 0 h state by AED = 0.63–0.95 between 12 and 48 h, whereas inoculated and mock samples collected at the same time differed by AED = 0.20–0.22. In healthy berries of a grapevine introgression line, its non-carrier sibling and ’Syrah’, sampled across development without pathogen challenge [38], AED between the first two sampled stages was 0.92–1.21 and reached 5.3–7.0 at late stages; this dataset differs in tissue and design and only shows how AED grows with the extent of transcriptional change. Mock samples are not a neutral baseline; time, handling and the mock treatment itself can change transcription. Changes in our susceptible control between time points partly overlap this range (0.35–0.90 vs 0.63–0.95). AED was not driven by a few genes: in this study, 3.3–6.8% of genes accounted for half of the total AED and 31.7–40.8% for 95% of it (21 comparisons; Additional file 5: Table S2). The external datasets showed similar distributions (half of the total from 2.8–6.9% of genes), so the AED values reflect shifts spread across many genes rather than a small set of outliers. Because AED depends on the gene set, normalization and design of each study, these values serve as reference magnitudes rather than thresholds. Synthetic per-gene noise of up to 10% of each gene’s standard deviation produced AED ≤ 0.008 (Additional file 4: Figure S3A).

Principal component analysis of all 26,169 genes reinforced the observed patterns (Figure 2 C/D). PC1 (40.3% variance) captured the dominant transcriptional gradient, with *Rpv12+1+3* showing the largest separation from the susceptible control, while PC2 separated *Rpv12, Rpv12+1 and Rpv12+1+3* from the susceptible genotype. PC3 further separated *Rpv12+1* from the other genotypes. Genes with the highest contribution to PC1 (Additional file 7: Figure S4; Additional file 8: Table S4) were enriched for proteins annotated with signal peptides (UniProt keyword ’Signal’; FDR = 3.35×10⁻⁷). Genes with the highest contribution to PC2 were enriched for domain annotations typical of NLR immune receptors (InterPro APAF-1 helical domain, FDR = 6.95×10⁻⁶; NB-ARC, FDR = 7.52×10⁻⁶); these annotations do not indicate cell-death activity. Top-contributing genes for PC3 were enriched for the UniProt keyword ’Hypersensitive response’ (three genes; FDR = 0.031), and those for PC4 for a mixed STRING local-network cluster that includes systemic-acquired-resistance annotations (three genes; FDR = 0.029). These are annotation enrichments and do not measure activation of the corresponding responses. In summary, all three resistant genotypes differed from the susceptible control already at inoculation, and they differed from each other mainly in how much they changed after inoculation.

### Genotype differences in the number of differentially expressed genes were largest at 6 hpi

A total of 9,459 unique genes (Additional file 9: Table S5) were differentially expressed across all experimental conditions (FDR < 0.05, |log₂FC| > 1). The distribution of DE genes among comparisons is summarized in Figure 3.

**Figure 3:**
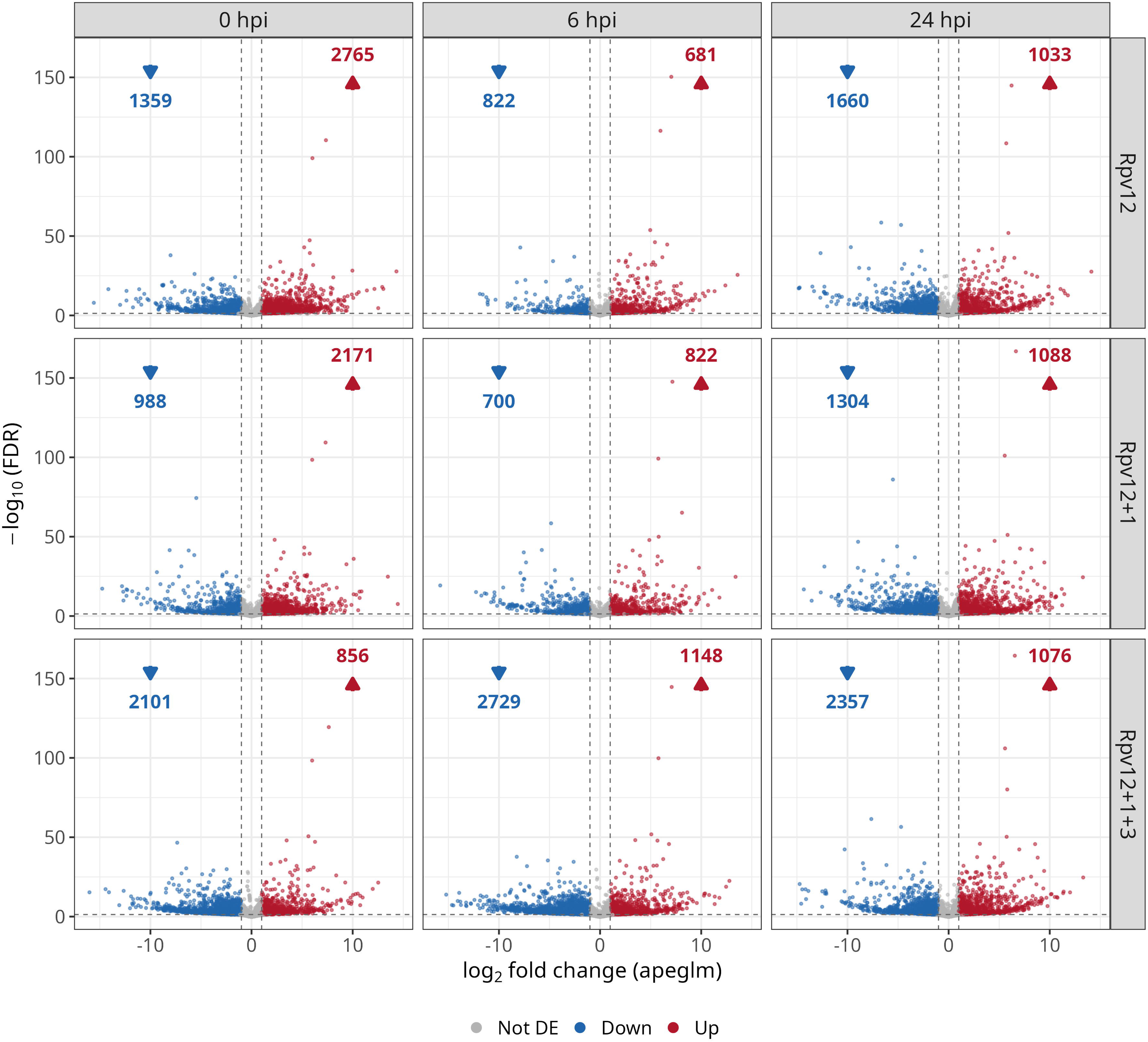
Volcano plots of differentially expressed genes for each genotype. *Rpv12*, *Rpv12+1*, *and Rpv12+1+3* (rows) are compared with the susceptible control at each time point (columns: 0, 6, 24 hpi). The x-axis shows log₂ fold change and the y-axis shows −log_10_(FDR). Red points indicate significantly upregulated genes (log₂FC > 1, FDR < 0.05), blue points indicate significantly downregulated genes (log₂FC < −1, FDR < 0.05), and gray points are not significant. Numbers in the upper corners show counts of significantly downregulated (↓, blue) and upregulated (↑, red) genes per condition.

We modeled these DEG counts and patterns in two complementary ways (Additional file 9: Table S5; Additional file 10: Table S6; Additional file 11: Figure S5). Models A-D test raw DEG counts per genotype and time point directly. Models E-F instead classify the same underlying DE calls by temporal-response category and by which genotypes share a given expression pattern, addressing a different question. Both are exploratory summaries: the counts and categories derive from the same samples and a shared susceptible reference, are not independent, and inherit the uncertainty of the threshold-based DE calls. Across resistant genotypes, DEG counts did not scale systematically with time point, genotype identity, or direction of change when considered in aggregate (Additional file 10: Table S6; Model A: time point FDR = 0.266, genotype FDR = 0.329, direction FDR = 0.360). DEG counts also did not scale with major resistance locus dosage (Model B: loci FDR = 0.380). Considered separately by genotype, none showed a significant trend in overall DEG count across time points (Model C: *Rpv12*: FDR = 0.691; *Rpv12+1*: FDR = 0.691; *Rpv12+1+3*: FDR = 0.691).

Per-time point models (Model D) showed that genotype was not associated with DEG count at 0 hpi (FDR = 0.639) but was at 6 hpi (FDR = 2.1 × 10⁻⁶) and 24 hpi (FDR = 0.015). In leave-one-replicate-out refits of the DESeq2 model (each of the 36 libraries omitted in turn), *Rpv12+1+3 had the most differentially expressed genes at 6 hpi in all 36 refits, with a median of 2.6 times as many as the higher of Rpv12 and Rpv12+1, whereas at 0 and 24 hpi the genotype with the most differentially expressed genes exceeded the second-ranked genotype by a median of only 1.3-fold (Additional file 10: Table S6).* Direction of change was not significantly skewed at 0 or 6 hpi (FDR = 0.492 and 0.138, respectively), whereas at 24 hpi downregulation was more common than upregulation (FDR = 6.3 × 10⁻⁶).

To describe these patterns further (Additional file 10: Table S6; Additional file 12: Table S7; Additional file 11: Figure S5), we classified each gene by its temporal-response category: IEV (initial variation; DE at 0 hpi, or at 0 and 6 hpi), ER (early response, at 6 and 24 hpi), TSR (transient stress response, at 6 hpi only), LR (late response, at 24 hpi only), SCh (sustained change, across all three time points), or CP (complex pattern, not fitting the above five), and by which genotypes share the identical pattern (pattern group: I, shared by all three genotypes; II, shared by *Rpv12* and *Rpv12+1*; III, shared by *Rpv12+1* and *Rpv12+1+3*; IV, shared by *Rpv12* and *Rpv12+1+3*; Va/Vb/Vc, specific to *Rpv12*/*Rpv12+1*/*Rpv12+1+3* respectively; VI, complex patterns spanning multiple genotypes). Gene numbers differed widely between pattern groups within every temporal-response category, and downregulated genes predominated among early-and late-response genes (Models E and F; Additional file 12: Table S7). Because these categories are built from the same DE calls, the model results are descriptive summaries rather than independent tests. Overall, genotype differences in the number of differentially expressed genes were concentrated at 6 hpi, and the counts did not follow the number of resistance loci.

### Comparison with an independently generated *Rpv12* transcriptome

To test whether aspects of the within-genotype temporal response recur in an independent *Rpv12*-carrying genotype, we compared our data against an independently generated dataset. Chitarrini et al. [29] profiled a distinct *Rpv12*-carrying genotype, derived from a cross between ’Bianca’ and the breeding line SK77-4/5 and unrelated to the material used here, following *P. viticola* inoculation of detached leaf discs (ENA study PRJEB28042). We reprocessed their raw reads (Additional file 13: Table S8) through the same PN40024.v4 alignment and quantification pipeline applied to our own libraries. After rlog transformation, sample-to-sample Pearson correlations separated libraries primarily by time after inoculation and secondarily by treatment (Additional file 14: Figure S6), suggesting that the reprocessed matrix retained the expected structure.

Because Chitarrini et al. [29] did not include a susceptible genotype, the only comparable axis is the within-*Rpv12* change over time after inoculation. We therefore compared our *Rpv12* 6 hpi and 24 hpi contrasts, each relative to 0 hpi, with their inoculated 12 h and 24 h samples, each relative to mock at 0 h, restricting both to the 24,338 genes tested in both studies. Chitarrini’s 12 h sampling is the nearest available proxy for our 6 hpi time point, not a match. Our temporal contrasts were taken from the full model including sequencing batch (∼batch + condition, all 36 libraries), because the *Rpv12* 0 hpi and 24 hpi libraries were sequenced in different batches.

At both time points, the two datasets shared more differentially expressed genes than expected by chance, and about three quarters of the shared genes changed in the same direction (Additional file 13: Table S8). At the early time point, 793 genes were shared (of 5,067 in our data and 2,452 in Chitarrini et al. [29]; 1.6-fold enrichment, one-sided hypergeometric p = 2.0 × 10⁻⁴⁵), of which 592 (74.7%) changed in the same direction, compared with 56.1% expected from the proportions of up-and downregulated genes in each study (one-sided Fisher’s exact test, p = 1.6 × 10⁻³²). At 24 hpi, 354 genes were shared (of 2,464 and 1,762; 2.0-fold, p = 6.4 × 10⁻³⁹), of which 261 (73.7%) agreed in direction (expected 59.9%; p = 2.0 × 10⁻¹¹). Because genes are not independent, these p-values should not be read literally. Because the same three plants per genotype were sampled at all time points, we refitted the model with the individual plant included; directional agreement was nearly identical (75.3% and 73.4%, expected 56.3% and 61.9%; Additional file 13: Table S8).

Gene ontology enrichment of the direction-concordant genes, based on their closest *Arabidopsis* homologs, did not return pathogen-recognition categories. Most concordant genes were transcribed at lower levels after inoculation in both studies (437 of 592 at the early time point and 213 of 261 at 24 hpi), and the enriched terms reflected this shared down-regulation: photosynthetic light harvesting at the early time point (FDR = 0.037), and lignin and phenylpropanoid catabolism (FDR = 6.5 × 10⁻⁵) and secondary cell-wall biogenesis (FDR = 2.2 × 10⁻⁴) at 24 hpi (Additional file 13: Table S8). In addition, genes that differed between *Rpv12* and the susceptible control at 0 hpi overlapped more than expected with genes that changed between 0 and 12 h in Chitarrini’s mock-treated samples (625 genes, 1.5-fold, p = 8.6 × 10⁻³⁰), including SOBIR1 and PAD4. We therefore interpret the shared temporal response largely as a stress and growth-adjustment program; the comparison does not establish an *Rpv12*-specific signature. Neither dataset includes a time-matched, unwounded control, so diurnal and handling effects may contribute to this shared signal.

### Co-transcriptional network density followed the magnitude of temporal change

To investigate how co-transcriptional organization differs among the examined genotypes, we constructed correlation-based co-transcriptional networks from 9,459 differentially expressed genes across genotypes and time points (Additional file 9: Table S5), using a Pearson correlation threshold of r ≥ 0.8077 (positive correlations only) (corresponding to the 99.5^th^ percentile of the correlation distribution; Additional file 15: Figure S7A).

For each gene of interest, a co-transcriptional module was defined as that gene together with every other gene correlated with it at r ≥ 0.8077 (positive correlations only), requiring at least three such correlated partners (four genes total) for a module to be reported. The modules were designed to define co-transcriptional neighborhoods around genes of interest as a candidate pool for structural interaction screening, under the assumption that physically interacting proteins are more likely to share transcriptional signatures than random gene pairs.

Network composition was stable across correlation thresholds: with all 9,459 differentially expressed genes as anchors, rebuilding the network at the top 1%, 0.5% and 0.1% of pairwise correlations (r ≥ 0.757, 0.8077 and 0.8839) gave 8,485, 7,299 and 4,252 modules with at least three partners (median 136, 72 and 31 partners), and module sizes shifted smoothly with the threshold (Additional file 15: Figure S7B; Additional file 16: Table S9). At all three thresholds, network edges were more often supported by STRING-db links than random pairs of a differentially expressed gene and any other gene, and the enrichment increased with stringency: 2.0-, 2.4- and 3.4-fold for all STRING evidence, 2.8-, 3.2- and 5.9-fold for experimental evidence only, and 2.6-, 3.1- and 4.9-fold when text-mining and curated-database evidence were excluded (Additional file 15: Figure S7C). Only a small fraction of edges (0.07–0.48%) had STRING support, as expected from STRING’s sparse coverage of grapevine. Edges of the primary network (rlog-transformed, condition-protected ComBat-corrected values; 938,485 edges among the 9,459 differentially expressed genes at r ≥ 0.8077) were also stable under a specified Gaussian noise model: the perturbed networks retained 99.6% of the noise-free edges at 0.5% noise and 95.3% at 5% noise (Jaccard similarity 0.993 and 0.929, respectively; 200 replicates per level). A separate sensitivity analysis on DESeq2 negative-binomial Pearson residuals, a different expression representation, gave similar values (99.4% and 93.1%).

Depending on score type, 13.7–17.5% of the modules at r ≥ 0.8077 (of those with STRING coverage) reached an empirical p < 0.05 for elevated STRING-db interaction or co-expression scores in a size-matched permutation test, compared with 5% expected for a single test; because modules overlap and are not independent, these proportions are reported descriptively. Module PC1 explained 85% of within-module variance on average (61% for genome-wide WGCNA modules), as expected for neighborhoods selected for strong correlation. These descriptive checks show that the modules are internally coherent, which their construction partly ensures, and partly consistent with external functional annotation; STRING associations include co-expression and transferred evidence and do not validate physical interactions in grapevine. Classical weighted gene co-expression network analysis (WGCNA) was evaluated as a comparator but was not used for candidate-pair selection, because it collapsed the transcriptome into a smaller number of very broad modules (the largest modules having 5,499 and 3,896 genes, respectively; Additional file 15: Figure S7D) that mix multiple distinct expression programs together.

At 0 hpi, the network had a density of 0.069 (mean degree 649) and a clustering coefficient of 0.68. Density declined from 0 hpi (density = 0.069) to 6 hpi (density = 0.041) and 24 hpi (density = 0.034); with three time points, no trend test is informative (Additional file 17: Figure S8). Across genotypes, *Rpv12* and *Rpv12+1* formed denser networks than the susceptible control (susceptible density = 0.053; *Rpv12* density = 0.101, *Rpv12+1* density = 0.091; susceptible mean degree = 498, *Rpv12* = 959, and *Rpv12+1 =* 859), whereas *Rpv12+1+3* showed reduced connectivity (density = 0.031; mean degree = 291). Density did not increase with the number of major resistance loci (Spearman ρ = −0.40; with four genotypes, such tests have little power). Because each network pools samples from several conditions, correlations largely reflect differences between condition means: per-genotype density followed the same order as the magnitude of within-genotype temporal change (mean AED; *Rpv12* > *Rpv12+1* > susceptible > *Rpv12+1+3*; Additional file 5: Table S2), so the two measures are not independent, and density differences do not by themselves indicate regulatory rewiring. A fixed threshold also corresponds to a different null exceedance rate at each sample size (for r ≥ 0.8077, about 0.4% with the nine samples of a per-genotype network and 0.07% with the twelve samples of a per-time-point network), so densities are compared only within each network type. Global clustering remained moderate (0.51-0.71) in all conditions. Network density therefore mirrors the magnitude of temporal change in each genotype and does not provide independent evidence of regulatory differences.

### A structural interaction screen based on the reference gene models

To examine whether proteins encoded by genes of interest could interact, they were screened against their co-transcriptional module members. We performed systematic AlphaFold2-Multimer screening on protein pairs selected from these co-transcriptional modules, from the plant-immunity literature and shared transcriptional patterns, and across modules (Additional file 18: Table S10). Because large-scale AlphaFold-Multimer interaction screens [44] show high sensitivity but limited specificity [45], and structurally similar repeat proteins could in principle yield spuriously plausible interfaces, each pair was evaluated across the five models produced by the five AlphaFold2-Multimer parameter sets from a shared MSA (ColabFold v1.6.2, AlphaFold2-Multimer v3) against a strict criterion: ipTM ≥ 0.75 in all five models, ≥5 interface contacts, and mean contact-filtered PAE < 10 Å across the five models. Predicted interactions were further required to survive a dedicated promiscuity screen (below) and, where applicable, to be distinguished from established literature-or homology-supported complexes reported separately as positive controls.

With the PN40024 v4 gene models, 10 pairs outside known complexes met all criteria (Additional file 18: Table S10, Figure 4A), alongside 44 pairs recovered within the known EDS1-SAG101-PAD4 complex family (positive controls; 32 of them EDS1– SAG101 pairs, ipTM 0.75–0.89, mean contact-filtered PAE 2.12–4.76 Å; Figure 4C), 10 pairs passing the interface criteria but flagged and excluded as promiscuous (see below), and 192 pairs with partial support that did not meet the full five-model threshold. Among these predictions, the SOBIR1-LRIP1 interaction, proposed a priori on the basis of homology to the tomato LeEIX2-SlSOBIR1 receptor complex, met the strict criterion: all five models had ipTM ≥ 0.75 (range 0.77-0.81, mean 0.79), and the mean contact-filtered PAE across the five models was 7.80 Å; four models were individually below 10 Å and one was at 18.7 Å, so the pair passes on the five-model mean rather than in every model. In all five models, the confident interface lay between the C-terminal end of the SOBIR1 LRR ectodomain (residues 185–209) and the N-terminal end of the LRIP1 gene model (residues 1–24), with 57–64 contacts per model, mean interface pLDDT of 85–90 (SOBIR1) and 82–87 (LRIP1) and mean contact PAE of 2.3–3.8 Å. The remaining contacts, between the cytoplasmic SOBIR1 kinase domain and LRIP1, had low confidence (mean PAE 20–26 Å) and account for the 18.7 Å model.

**Figure 4:**
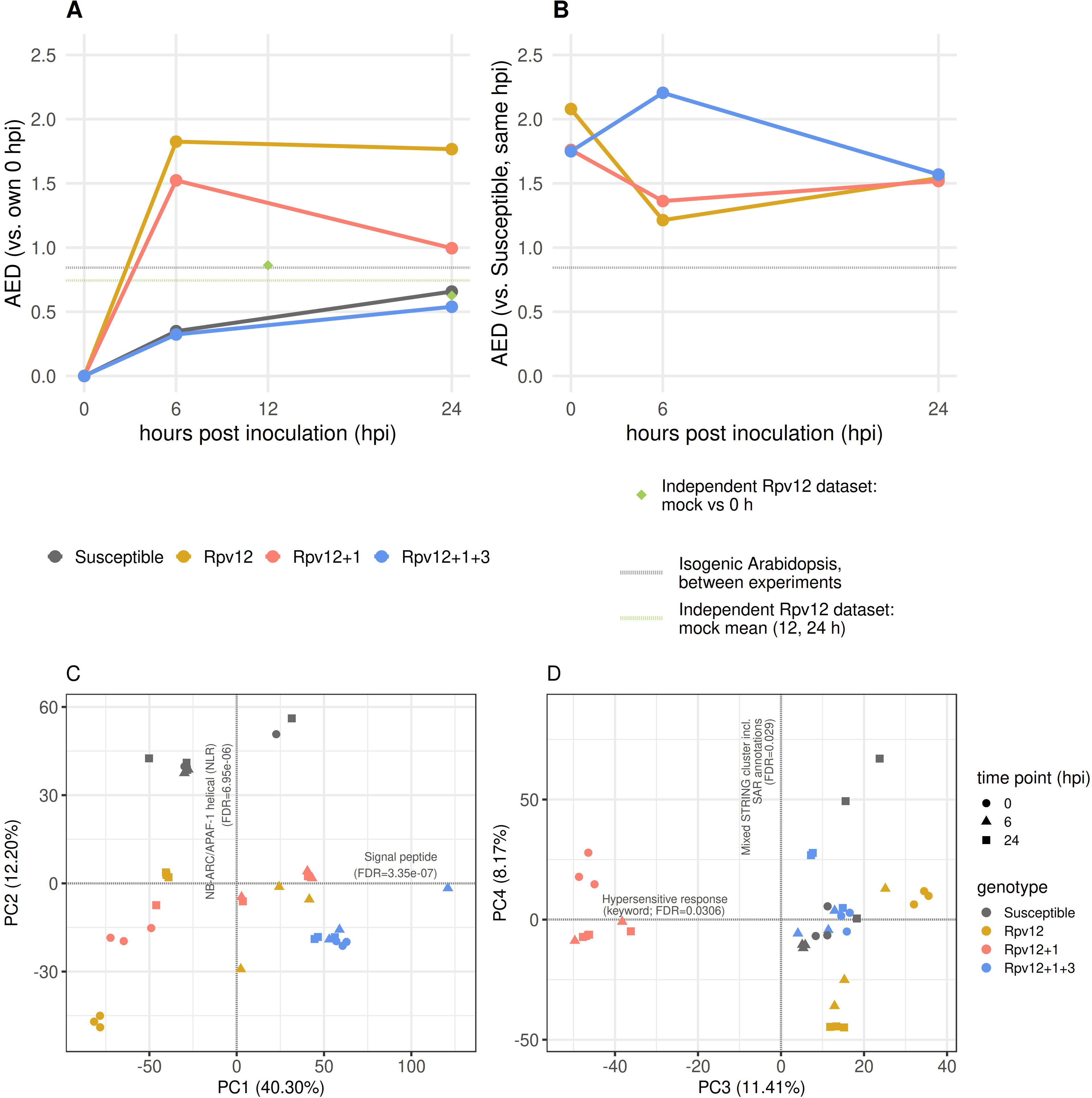
AlphaFold2-Multimer interaction predictions based on PN40024 v4 gene models. (A) The nine-gene, ten-edge AlphaFold2-Multimer network anchored on SOBIR1 (Vitvi17g00964) and LRIP1 (Vitvi09g04475); BAK1-like = Vitvi12g01277, a 97-residue BAK1 fragment in the v4 annotation. All edges meet the strict five-model criterion (ipTM ≥ 0.75 in all five models, ≥ 5 interface contacts, mean contact-filtered PAE < 10 Å). Edge labels give the mean contact-filtered PAE (Å) and edge color its class (green, < 3 Å; amber, 3–5 Å; orange, 5–10 Å); the ten means range from 1.68 to 9.91 Å, and single models can exceed 10 Å (one SOBIR1–LRIP1 model, 18.7 Å). Line type gives the source of the pair (Additional file 18: Table S10): solid, within a co-transcriptional module; dashed, cross-module pair; dotted, receptor-family screen of BRI1, BAK1, CLV1 and BAM1 homologs and leucine-rich repeat genes of the SOBIR1 and LRIP1 modules. Dollar signs mark genes with significantly increased transcript abundance in *Rpv12* relative to the susceptible control at 0 hpi (DESeq2, FDR < 0.05, | log₂FC| > 1); 8 of 9 genes, MRH1 excepted. ‡, gene model shorter than its closest *Arabidopsis* homolog (more than 25 residues of the homolog missing at either end; see Discussion, Limitations). None of the ten edges met the strict criterion when the pairs were predicted again with longer gene models from newer annotations (Additional file 18: Table S10, sheet AF2_rerun_gene_models), so the edges depend on the v4 gene models; interactions are predicted, not experimentally validated. (B) Schematic comparison with two characterized SOBIR1-dependent systems: the tomato system, in which EIX binds both LeEIX1 and LeEIX2 but only LeEIX2 signals and associates with SlSOBIR1 [24, 55], and BAK1 binds the decoy receptor LeEIX1, which forms an EIX-induced heterodimer with LeEIX2 (dashed line) and attenuates its signaling [55], and the *Arabidopsis* LRIP1–SOBIR1–BAK1 system, in which BAK1 is recruited in a pathogen-specific, elicitor-dependent manner [46]. SOBIR1 is drawn with extracellular LRRs, a transmembrane segment (TM) and a cytoplasmic kinase domain; *Arabidopsis* LRIP1 is an LRR-only protein. (C) EDS1 (Vitvi17g04216) and its seven predicted interaction partners: five SAG101-like paralogs (SAG101-I = Vitvi05g00062, SAG101-II = Vitvi14g03030, SAG101-III = Vitvi14g03031, SAG101-IV = Vitvi14g03032, SAG101-V = Vitvi14g04642), one divergent paralog retaining only the Lipase_3 domain (SAG101-like† = Vitvi14g03033), and PAD4 (Vitvi07g01908); ‡ as in panel A. All pass the strict five-model criterion (mean ipTM 0.77–0.88, mean PAE 2.12–4.76 Å); EDS1–SAG101 and EDS1–PAD4 pairs in which both gene models are complete also passed (Additional file 18: Table S10). (D) Pfam-A domains retained after strict filtering (genome-wide HMMER v3.3.2 run; see Methods and Additional file 18: Table S10) for the 17 genes in panels A and C, by domain category; a gene can appear in more than one category. (E) Promiscuity panel: each anchor was screened against a fixed panel of 100 candidate proteins (80 receptor-family, 20 non-receptor) under the strict criterion; SOBIR1 passed with 0 of 100 and LRIP1 with 2 of 94 candidates with complete model coverage (2.1%). For comparison, 0 of 366 pairs in the pre-specified transcription-factor negative-control set passed. These are computational checks, not experimental validation.

To test whether generic AlphaFold2-Multimer "stickiness" could explain these predictions (Additional file 18: Table S10, Figure 4E), SOBIR1 and LRIP1 were each screened against a fixed, controlled panel of 100 candidate proteins (80 receptor-kinase-family genes and 20 non-receptor background genes; 200 pairwise predictions for the two anchors, within a 1,600-pair screen of 16 query proteins). Under the strict five-model criterion, SOBIR1 passed against 0/100 panel candidates and LRIP1 against 2/94 candidates with complete model coverage (2.1%), indicating that neither protein forms high-confidence interfaces indiscriminately within this panel (Additional file 18: Table S10, Figure 4E).

The behavior of the screen was checked with a single pre-specified negative-control set defined before screening: four transcription-factor hubs (PRE6, WRKY51, WRKY55, NAC29) and their co-expression partners (422 declared pairs; 366 with complete five-model coverage). None of these 366 pairs met the strict criterion (0%); the remaining 56 pairs lacked recoverable AlphaFold2 output and are reported as unpredicted rather than negative. Cross-module pairings among immunity-related proteins from different co-transcriptional modules (n=325) are reported descriptively as part of the discovery screen rather than as an independent specificity control, since several of the compared hub proteins, including SOBIR1 and LRIP1 themselves, were candidates under active investigation rather than a pre-specified negative set; two of these 325 pairs met the strict criterion (0.6%), one being the SOBIR1-LRIP1 pair discussed above and the second a further predicted interaction between LRIP1 and MRH1 (Vitvi19g00098). With the v4 gene models, the screen therefore gave ten candidate pairs that passed all computational checks; whether they depend on the gene models is examined below.

### Receptor-related genes with elevated transcripts in *Rpv12* at 0 hpi and their predicted interactions

Under the strict five-model criterion, SOBIR1 (Vitvi17g00964), an adaptor kinase for RLP-type receptors, has three predicted partners: Vitvi09g04548 (LRIP1-like; mean ipTM 0.82, range 0.81-0.83; mean contact-filtered PAE 9.91 Å), Vitvi09g01951 (RGI1; mean ipTM 0.80, range 0.79-0.82; mean PAE 8.35 Å), and LRIP1 (Vitvi09g04475; scores given above) (Figure 4A). The grapevine LRIP1 homolog has four predicted partners: RLP34 (Vitvi01g04419; mean ipTM 0.82, mean PAE 2.33 Å), an LRIP1-like protein (Vitvi01g04416; mean ipTM 0.77, mean PAE 3.95 Å), MRH1 (Vitvi19g00098; mean ipTM 0.83, mean PAE 8.92 Å) and SOBIR1. Within this nine-gene network, LRIP1 and Vitvi01g04416 are the highest-degree nodes (four predicted partners each), with SOBIR1 and Vitvi09g04548 as secondary hubs (three each). Beyond this nine-gene network, additional protein pairs meeting the single-model interface criterion in at least one of the five models are reported in Additional file 18: Table S10 and together define the 43-gene set considered in the domain-architecture analysis below.

Of SOBIR1’s two non-LRIP1 partners, only Vitvi09g04548 met the differential-expression criterion for upregulation at 0 hours post-inoculation in *Rpv12* but in neither multilocus genotype (log2FC=+3.91, padj=1.0 × 10⁻⁴), whereas in *Rpv12+1* (log2FC=+0.36) and *Rpv12+1+3* (log2FC=-0.78) it did not meet the differential-expression criterion relative to the susceptible control. Vitvi09g01951 (RGI1) does not fit this pattern: also upregulated in *Rpv12* at 0 hpi (log2FC=+2.75, padj=0.0041), but significantly downregulated in *Rpv12+1+3* (log2FC=-3.19, padj=0.0097), indicating genotype-specific regulation. Direct comparisons between the resistant genotypes at 0 hpi (Additional file 19: Table S11) gave a consistent picture for the partners of the two hubs: Vitvi01g04416, Vitvi09g04548 and RLP34 had higher transcript levels in *Rpv12* than in both *Rpv12+1* (log2FC = +1.46 to +2.72) and *Rpv12+1+3* (log2FC = +3.84 to +6.36), as did LRIP1 (+1.36 and +3.23), whereas SOBIR1 and the BAK1-like gene were higher than in *Rpv12+1+3* (+2.50 and +1.06) but did not meet the fold-change criterion against *Rpv12+1* (+1.00 and +0.84). Thus, at 0 hpi eight of these nine receptor-related genes had higher transcript levels in *Rpv12* than in the susceptible control, and four of them (Vitvi01g04416, Vitvi09g04548, RLP34 and LRIP1) were also higher than in both multilocus genotypes.

The two hubs were connected directly (SOBIR1–LRIP1) and indirectly through Vitvi01g04416 and Vitvi09g04548, each of which passed the strict criterion with one hub but not the other. These connections were predicted from the v4 gene models and, as described below, were not reproduced with longer models.

Pfam domain annotation of the 43 genes in the structural interaction set (HMMER; Additional file 18: Table S10) retained at least one domain in 41 genes after strict filtering (see Methods). Leucine-rich repeat domains were present in 22 of these 41 genes (15 of the 32 genes in pairs passing the strict criterion), EDS1/PAD4/SAG101-family domains (EDS1_EP, Lipase_3) in 14 (14), and protein kinase domains in 8 (6). Kinase domains occurred only in homologs of receptor-like kinases; several of these homologs, including RGI1, GSO2 and the BAK1-like protein, lack a kinase domain in their PN40024 v4 gene models.

### The predicted receptor interactions depended on the gene models

Because seven of the nine genes in the predicted LRIP1/SOBIR1 network have PN40024 v4 models shorter than their *Arabidopsis* homologs (Figure 4A), we compared them with models of the same loci in newer annotations of the PN40024 genome (Additional file 18: Table S10). These annotations contain complete models for four of the seven shortened genes, including LRIP1 with its signal peptide. The longer models are not simple extensions of the v4 models: the genome sequence at these loci was nearly identical between assemblies (99.8–100%; 97.5% for Vitvi01g04416, which lies in a duplicated segment), but for five of the seven genes the longer and v4 proteins shared only 55–87% identity, because the annotations place exons and reading frames differently (sheet gene_models_loci). Two of the complete models are also uncertain: the 1,053-residue LRIP1 model is 16% longer than its homolog and lies in a cluster of LRIP1-like genes, and the 620-residue BAK1-like model has the 11-exon structure of *Arabidopsis* BAK1 but spans 35 kb (3.6 kb in *Arabidopsis*), and its last three introns are seen only in multi-mapped RNA-seq reads. Across the 42 genes of the structural interaction set with an *Arabidopsis* homolog, the number of complete models ranged from 12 to 24 per annotation (17 in v4), and 14 genes, including EDS1 (Vitvi17g04216), GSO2 and RGI1, lacked a complete model in all four (sheet gene_models_43). Predicted again with the longer models, none of the ten pairs met the strict criterion (sheet AF2_rerun_gene_models): eight showed no consistent interface (ipTM 0.20– 0.54; mean contact-filtered PAE 17–29 Å), and two SOBIR1 pairs were weakened (two of five RGI1–SOBIR1 models with ipTM 0.68 and contact-filtered PAE 6 Å; one Vitvi09g04548–SOBIR1 model meeting the single-model criterion). By contrast, all 16 EDS1–SAG101 and EDS1–PAD4 pairs with complete v4 models passed the strict criterion in the screen (ipTM 0.82–0.89; mean PAE 2.3–2.9 Å). Because neither set of models has been validated, these comparisons do not show which predictions are correct.

### Predicted EDS1–SAG101–PAD4 paralog network

Under the strict five-model criterion, EDS1 (Vitvi17g04216) had seven predicted interaction partners: five SAG101-like grapevine paralogs (Vitvi05g00062, Vitvi14g03030, Vitvi14g03031, Vitvi14g03032, Vitvi14g04642; mean ipTM 0.77-0.88, mean contact-filtered PAE 2.1-4.8 Å), a more divergent paralog retaining only the Lipase_3 domain (Vitvi14g03033; mean ipTM 0.84, mean PAE 2.7 Å), and PAD4 (Vitvi07g01908; mean ipTM 0.88, mean PAE 2.8 Å), EDS1’s other canonical *Arabidopsis* complex partner [23]. Two further SAG101-like paralogs (Vitvi14g04643, Vitvi14g04644) showed partial evidence only (mean ipTM 0.67-0.76) and did not meet the strict criterion (Additional file 18: Table S10). Vitvi14g03033 has a 332-residue model in all three annotations that contain it and was also predicted to bind each of the four EDS1 paralogs with complete models (ipTM 0.81–0.87), so this prediction probably reflects docking of its lipase-like domain rather than a specific partnership (Additional file 18: Table S10, sheets gene_models_43 and positive_control_EDS1_SAG101_PA).

Six of these seven predicted interaction partners (Figure 4C), together with EDS1 itself, were also transcriptionally upregulated in *Rpv12* at 0 hpi (canonical DESeq2, FDR<0.05 and log2FC>1): EDS1 (log2FC=+2.69), PAD4 (log2FC=+1.11), four of the five SAG101 paralogs (Vitvi05g00062 log2FC=+1.29, Vitvi14g03030 log2FC=+1.40, Vitvi14g03031 log2FC=+2.08, Vitvi14g03032 log2FC=+3.53), and the divergent Lipase_3-only paralog (Vitvi14g03033, log2FC=+4.91). EDS1 and three of these SAG101 paralogs (Vitvi05g00062, Vitvi14g03031, Vitvi14g03032) were also elevated, less strongly, in *Rpv12+1* (log2FC=+1.13 to +1.71), whereas none of these genes was elevated in *Rpv12+1+3*. Among the helper NLRs carrying the RPW8 domain (Additional file 9: Table S5, sheet helper_NLRs_RPW8), three of the eleven NRG1 homologs (Vitvi07g01315, Vitvi10g01495, Vitvi10g02048) had higher transcript levels at 0 hpi in *Rpv12* (log2FC = +2.75 to +3.29) and also in *Rpv12+1+3* (Vitvi07g01315) or *Rpv12+1* (Vitvi10g01495, Vitvi10g02048), whereas the single ADR1 homolog (Vitvi18g00850) was not differentially expressed. One predicted interaction partner, Vitvi14g04642 (mean ipTM 0.86), showed only borderline upregulation at this time point (log2FC=+1.02, padj=0.077). Thus, EDS1-family transcripts were elevated mainly in *Rpv12* at inoculation, and predicted EDS1 complexes involving complete gene models agree with known complexes.

### Functional associations among immunity-related genes elevated in *Rpv12*

Among the *Arabidopsis* homologs of the proteins in these two predicted networks (ten proteins, because several grapevine paralogs share one homolog), STRING-db (all evidence channels, including text mining and curated databases) connected six (EDS1, PAD4, SAG101, SOBIR1, BAK1 and LRIP1), reflecting their known functions. *Arabidopsis* LRIP1 (AT2G34930) is an LRR protein that associates with SOBIR1 [46] (Figure 4B; see Discussion).

Extending this analysis to the broader set of immunity-related genes established in the literature [47–52] and transcribed differently in *Rpv12* than in the susceptible control at 0 hpi (18 genes total: EDS1, SAG101, PAD4, SOBIR1, BAK1-like protein, MIK2-like (Vitvi07g02294), LRIP1, LAZ5, SARD4, WRKY51, WRKY55, MRH1, RLP34, GSO2, FER, RGI1, Vitvi12g02607/AT5G17680/ MVA3.30, and ACD6; 17 of the 18 at FDR < 0.05 and |log2FC| > 1, 15 at higher and LAZ5 and FER at lower levels, MRH1 not significant; eight of the 17 met the criterion only in *Rpv12* among the resistant genotypes; Figure 5A) were connected by 22 STRING-db edges (*Arabidopsis* homologs, as above), forming a single, densely connected 13-gene component (EDS1, PAD4, SAG101, BAK1-like protein, SOBIR1, MIK2-like, LRIP1/AT2G34930/ F19I3.16, LAZ5, SARD4, WRKY51, RLP34, ACD6, and MRH1) alongside five genes without STRING-recognized connections within this set (WRKY55, GSO2, FER, RGI1, Vitvi12g02607/AT5G17680/ MVA3.30). Because these genes were selected from the immunity literature, these connections and their annotations (for example ‘Systemic acquired resistance’ and ‘Positive regulation of defense response’; Figure 5B) are expected and summarize known functions of the homologs rather than validating the grapevine results. In a phased genome analysis of an *Rpv12*-carrying accession, three ACD6-like genes lie within a differentially expressed gene cluster at the *Rpv12* locus [53].

**Figure 5:**
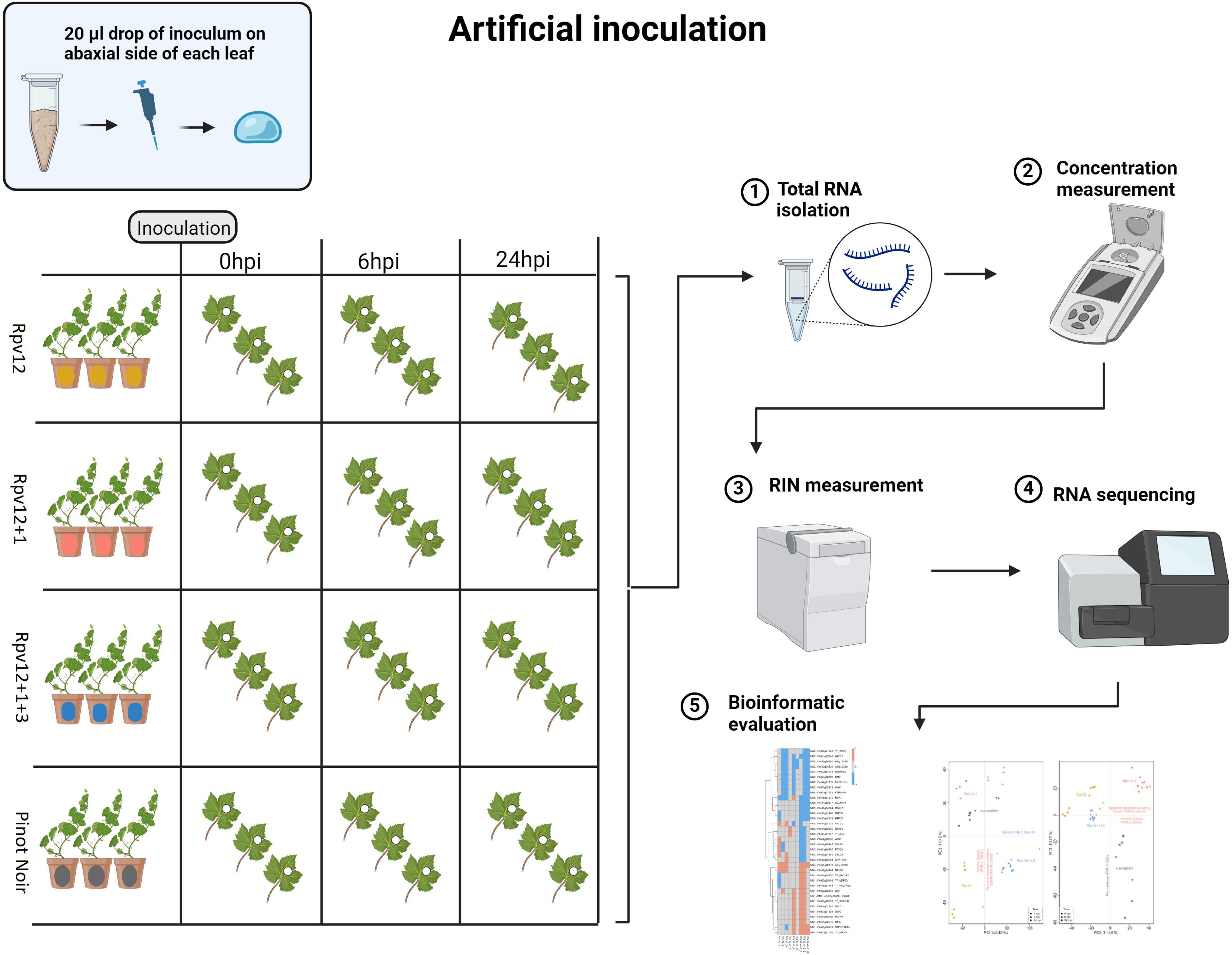
Transcript levels and STRING functional associations of 18 immunity-related genes in the resistant genotypes. (A) Log2 fold-change heatmap for 18 immunity-related genes across three genotypes carrying different resistance-locus combinations (*Rpv12*, *Rpv12+1*, *Rpv12+1+3*) at 0, 6, and 24 hpi relative to the susceptible reference (DESeq2 apeglm log2FC). Genes are grouped by network (colored bar, right): LRIP1/SOBIR1 group (blue; receptor-related genes, mostly from Figure 4A, and MIK2-like), EDS1–SAG101–PAD4 group (orange), and other literature-established immunity-related genes (green). Asterisks mark differential expression (DESeq2, FDR < 0.05 and |log2FC| > 1), as reported in Additional file 9: Table S5. BAK1-like = Vitvi12g01277, a 97-residue BAK1 fragment in the v4 annotation; SAG101 is represented by the divergent paralog Vitvi14g03033 (Lipase_3 domain only); MIK2-like = Vitvi07g02294; RGI1 = Vitvi09g01951. (B) STRING-db functional-association network of the same 18 genes (combined score ≥ 0.4). Edge width and color intensity scale with STRING combined score. Grapevine genes are represented by their *Arabidopsis* homologs; edges include all STRING evidence channels, including text mining. Node fill indicates network group (see legend); log2 fold changes are shown in panels A and C. Five genes (FER, GSO2, MVA3.30, RGI1, WRKY55) show no STRING-recognized interaction within this gene set. (C) Summary table of the 18 genes: network group, *Rpv12* log2 fold-change at 0 hpi, and STRING connectivity status.

## DISCUSSION

Across the four genotypes, the main differences lay in the transcriptional state at inoculation and in the extent of change afterwards: all three resistant genotypes differed from the susceptible control already at 0 hpi, genotype differences in the number of differentially expressed genes were largest at 6 hpi, and *Rpv12+1+3* changed least relative to its own 0 hpi state. The temporal response shared with an independent *Rpv12*-carrying dataset was dominated by general stress-and growth-related genes. In the *Rpv12* genotype, receptor-related and EDS1-family transcripts were elevated at inoculation, whereas the receptor interactions predicted from the reference gene models did not hold when the models changed. The sections below discuss these results in turn, followed by the limitations of the design.

### Transcriptional organization differed among the examined resistance genotypes

Transcriptional patterns differ among the three resistant genotypes; with one genotype per locus combination, these differences cannot be attributed to the number of loci.

Instead, each genotype exhibits distinct transcriptional patterns: *Rpv12 had higher transcript levels of LRIP1/SOBIR1-related receptor genes and EDS1–SAG101–PAD4-family genes at inoculation; Rpv12+1 moved away from its 0 hpi state by 6 hpi and partly back by 24 hpi;* and *Rpv12+1+3* changed least relative to its own 0 hpi state, with a correspondingly sparser genome-wide co-expression network, although it carries the largest combination of resistance loci. These differences are compatible with field-level evidence that locus pyramiding in grapevine does not straightforwardly sum the properties of its component loci: defeated or partially compromised resistance loci have been proposed to act as "Trojan horses" that can compromise, rather than reinforce, resistance pyramiding [32], and independent field surveys have already documented *Plasmopara viticola* strains overcoming pyramided *Rpv1-Rpv3.1* combinations in France [5], and strains overcoming *Rpv12* itself have also been reported [30].

The transcriptional differences observed in *Rpv12* at inoculation (0 hpi) need cautious interpretation: the independent *Rpv12*-carrying dataset (Chitarrini et al. [29], reprocessed for this study) cannot test the 0 hpi difference directly, because it lacks a susceptible genotype, but the genes that distinguish our *Rpv12* genotype at 0 hpi, including SOBIR1 and PAD4, are enriched among genes that change over time in its mock-treated samples. Part of this signature may therefore involve transcripts that respond to time of day or handling, and the temporal response shared by the two *Rpv12*-carrying genotypes is dominated by general stress-and growth-related functions rather than by a resistance-specific signal unique to any one pyramided genotype. The relationship between these transcriptional patterns and resistance remains unknown.

Field observations suggest that multilocus genotypes show effective resistance, but this cannot substitute for controlled phenotypic quantification, and the documented breakdown of other pyramided *Rpv* combinations in the field [5] is a further reason for caution before assuming that transcriptional differences alone predict durable resistance. Distinguishing these alternatives requires paired transcriptomic and resistance assays measuring pathogen growth, sporulation, and disease severity.

### Predicted receptor interactions depended on the gene models

With the v4 gene models, the predicted LRIP1/SOBIR1 interactions resembled established SOBIR1-dependent receptor systems. In tomato, the RLP LeEIX2 recognizes fungal xylanase [54] and associates with SOBIR1 [24], whereas its paralog LeEIX1 acts as a decoy receptor that attenuates this response [55, 56]. The ELR-SOBIR1 complex of *Solanum microdontum* recognizes *Phytophthora infestans* elicitins and functions as a two-component receptor [57]. The *Arabidopsis* LRR-only protein LRIP1 associates constitutively with SOBIR1 and recruits BAK1 in an elicitor-dependent manner, with allelic variants conferring resistance to either *Pseudomonas syringae* (via salicylic acid signaling) or *Botrytis cinerea* (via camalexin biosynthesis) [46]. In these systems a single adaptor serves sensors of unrelated ligands: specificity resides in the RLP or LRR-only sensor, and signaling competence in SOBIR1 and the recruited SERK co-receptor BAK1 [24, 25, 46, 57]. However, none of the predicted receptor interactions was reproduced with longer gene models from newer annotations, whereas EDS1 complexes with complete models were predicted consistently (see Results). The resemblance is therefore architectural, and the predictions do not indicate that a comparable SOBIR1-dependent module operates in grapevine. The promiscuity panel and the negative-control set argued against generic AlphaFold2-Multimer "stickiness" [45], but they could not detect this dependence on the gene models.

In the v4-based network, two LRIP1 paralogs (Vitvi01g04416 and Vitvi09g04548) were predicted to interact with Vitvi12g01277, a 97-residue BAK1-like fragment; with the 620-residue BAK1 model of a newer annotation, these interfaces were not predicted. In tomato, attenuation of EIX signaling by the LeEIX1 decoy requires BAK1 [55], in *Arabidopsis* RLP30-mediated immunity to necrotrophic fungi requires both SOBIR1 and BAK1 [58], and SOBIR1 is required for cell death triggered by perturbation of BIR1, an RLK that negatively regulates multiple RLP-mediated resistance pathways [22].

Whether a SERK protein associates with grapevine LRIP1 or SOBIR1 therefore remains open. Overall, the structural screen yielded hypotheses whose basis depends on gene annotation rather than candidate interactions ready for experimental testing.

### Elevated receptor-related and EDS1-family transcripts in *Rpv12*: a PTI–ETI hypothesis

At the time of inoculation, before pathogen penetration, SOBIR1, an adaptor kinase for RLP-type pattern recognition receptors, and EDS1 both had higher transcript levels in the *Rpv12* plants. EDS1 has recently been shown to act as a signaling hub capable of activating both PTI and ETI outputs [23], and forms functional heterodimers with PAD4 or SAG101 that orchestrate distinct immunity branches: EDS1-SAG101 operates with NRG1 in TNL-mediated ETI and cell death, while EDS1-PAD4-ADR1 modulates transcriptional defenses in basal immunity and reinforces CNL-triggered responses [17, 23]; the relative contribution of the ADR1 and NRG1 helper-NLR subfamilies to these outputs is itself species-variable rather than fixed [59]. In *Arabidopsis*, SOBIR1 associates in a ligand-independent manner with EDS1, PAD4 and ADR1-family helper NLRs, linking the receptor-like protein RLP23 to the EDS1–PAD4–ADR1 node and suggesting a preformed, SOBIR1-anchored signaling platform at the plasma membrane [23]. The co-elevation of SOBIR1, LRIP1-related and EDS1-family transcripts in the *Rpv12* genotype is consistent with such a connection, although whether grapevine SOBIR1 engages EDS1–PAD4 in this way remains to be tested. In *Rpv12*, transcripts of SAG101-like paralogs and of NRG1 homologs, the helper partners of EDS1–SAG101 in TNL-triggered signaling [16], were both higher at inoculation.

In grapevine, however, the EDS1 family shows structural divergence from *Arabidopsis*: a naturally occurring VvEDS1 splicing variant encoding only the N-terminal domain (VvEDS1^Nterm^, arising from retention of an intron containing a premature stop codon) forms stable homodimers that are structurally incompatible with PAD4 heterodimer formation, though the physiological relevance of this truncated isoform itself remains unconfirmed by the original study [60]; *Vitis* EDS1, by contrast, was separately shown by BiFC to physically interact with both *Vitis* and *Arabidopsis* PAD4 [61], and our AlphaFold2-Multimer screen also predicts a grapevine EDS1–PAD4 interaction (ipTM 0.88).

VvPAD4 is additionally unable to functionally substitute AtPAD4 in complementation assays [61]. A predicted EDS1 interaction partner in our dataset, a SAG101-like protein retaining only the Lipase_3 domain but lacking the EP domain required for canonical EDS1-SAG101 signaling (SAG101-like; UniProt F6HV27; ∼40% identity to SAG101), is upregulated alongside EDS1, though its signaling capacity remains unclear; because it was predicted to bind every EDS1 paralog with a complete model, this prediction does not indicate a specific partner. Four canonical SAG101 paralogs meet the upregulation threshold (FDR<0.05). Although the *Rpv12* locus has been reported to contain CNL candidate genes [12], which are generally considered EDS1-independent, *Rpv12 plants showed higher transcript levels of EDS1, SOBIR1 and LRIP1-related receptor genes, which is compatible with, but does not demonstrate, PTI–ETI co-activation [19, 62]; pathway activity was not measured.* Transcript abundance does not necessarily reflect protein levels or functional activity, and genetic perturbation and interaction assays would test whether SOBIR1 and EDS1 are required for resistance.

### Limitations and future directions

The main limitation of this study is also its central message: the unit of comparison is the genotype, not the resistance locus. Each locus combination is represented by a single breeding genotype rather than by near-isogenic material, so the effects of the *Rpv* loci cannot be separated from those of the genetic background into which they were introgressed. Because all three resistant genotypes derive from crosses involving ’Pinot Noir’, the susceptible control shares part of their genetic background; this narrows, but does not remove, the background differences underlying the comparisons with the control. All three resistant genotypes already differed from the susceptible control at 0 hpi, before pathogen penetration, and *Rpv12+1+3* changed less relative to its own 0 hpi state than *Rpv12* and *Rpv12+1* (see Results). The elevation of LRIP1/SOBIR1-related and EDS1–SAG101–PAD4-family transcripts should therefore be read as a property of the *Rpv12* genotype examined here rather than of the *Rpv12* locus. In *Rpv12+1*, only some of these transcripts (LRIP1, EDS1 and three SAG101 paralogs) met the differential-expression criterion, with smaller fold changes, and in *Rpv12+1+3* none was upregulated, although both genotypes also carry *Rpv12*. Compared directly, *Rpv12* had higher transcript levels than *Rpv12+1+3* for ten of these eleven genes and than *Rpv12+1* for six (Additional file 19: Table S11). This pattern argues against a simple locus effect, and whether it reflects effects of the additional loci or differences in genetic background cannot be resolved without isogenic single- and multilocus lines. This context dependence agrees with bioassay evidence that the same pyramided loci can have beneficial, neutral or detrimental effects on resistance depending on the locus combination, the genetic background and the pathogen strain [32]. For the same reason, resistance phenotypes reported for these locus combinations in other breeding populations cannot be assumed for the lines studied here, and, apart from the breeders’ field ratings (see Methods), no disease phenotypes were quantified on this material. Reads were also mapped to the PN40024.v4 reference rather than to haplotype-resolved assemblies of the introgressed regions; long-read sequencing would allow allele-specific transcript analysis and checks for reference-mapping bias at the *Rpv* loci and in the surrounding background.

All findings are based on transcript abundance and do not address protein abundance, post-translational modification or functional activity. Proteomics and functional assays (ROS production, callose deposition, MAPK activation) are required to test whether the transcriptional differences translate into altered immune capacity. In characterized RLP/LRR-only–SOBIR1–BAK1 systems, SOBIR1 and BAK1 kinase activities are required for immune signaling [25, 46]; the structural predictions do not show that the grapevine proteins interact or that such interactions contribute to resistance. The receptor predictions were not robust to the gene models used, and the structural models use PN40024 reference sequences rather than those of the genotypes studied. Testing these hypotheses requires verified gene models, followed by interaction assays (for example co-immunoprecipitation or split-reporter assays in *Nicotiana benthamiana*) and genetic perturbation. The structural thresholds used (ipTM ≥ 0.75 in all five models, ≥ 5 interface contacts, mean contact-filtered PAE < 10 Å) fall within the ipTM range of 0.6– 0.8 that the AlphaFold 3 documentation describes as a "gray zone where predictions could be correct or incorrect", with values above 0.8 indicating confident predictions [63]; these values are confidence guidelines, not a validated interaction classifier. No single threshold is universally validated, and recent benchmarks show that no single score generalises across all complexes [64, 65]; PAE-derived contact probabilities, which underlie our contact-filtered PAE criterion, have been shown to be well calibrated to the fraction of native interface contacts in solved structures [66]. Our criterion is therefore an operational screening rule that does not rely on ipTM alone, but on its combination with consistency across five models, interface-contact and PAE filters, the promiscuity panel, and the negative- and positive-control sets, which are computational checks rather than experimental validation.

Structural predictions used PN40024 v4 gene models, which are computational predictions [36]. For seven of the nine genes in the predicted LRIP1/SOBIR1 network, and for EDS1 and three of its predicted partners, the gene model is shorter than its closest *Arabidopsis* homolog (marked in Figure 4A, C): the LRIP1 model corresponds to residues ∼363–770 of its 905-residue homolog and lacks the N-terminal region including the signal peptide, RGI1 and GSO2 lack a kinase domain, and the BAK1-like gene model is a 97-residue fragment. When the ten candidate pairs were predicted again with the longer models of newer annotations, none met the strict criterion, whereas EDS1 complexes with complete models did, so these predictions depend on the gene models used. The longer models are themselves unverified, and two of them are uncertain (see Results), so the comparison does not show which prediction is correct. Discrepancies between gene models of leucine-rich repeat receptors are a known problem at such clustered, repetitive loci: three public annotations of the rice Nipponbare genome disagreed extensively for these genes, which led to their manual curation [67]. The grapevine loci examined here show the same pattern, with nearly identical genome sequence but different exon structures across annotations. However, a locus can also encode several protein isoforms through alternative splicing, which is widespread in grapevine [68] and explains many of the differences between cloned transcripts and gene models of leucine-rich repeat receptor kinases [69]; which of the predicted proteins are produced therefore needs to be verified, for example by long-read transcript sequencing [70] or proteomics [71]. More generally, AlphaFold2 predictions combined with checks of domain architecture against close homologs could serve as a quality assessment of gene models for tailored gene sets, alongside BUSCO completeness scores [72], or as a training signal for predictors of coding sequences.

The temporal dynamics do not resolve into a simple early-versus-late contrast between genotypes. Genotype effects on the number of differentially expressed genes were concentrated at 6 hpi across the resistant genotypes jointly (Additional file 10: Table S6, Model D), whereas the LRIP1/SOBIR1- and EDS1–SAG101–PAD4-family transcripts were already elevated in the *Rpv12* genotype at 0 hpi. The 6 hpi difference was consistent across leave-one-replicate-out refits but was not re-tested under alternative thresholds for calling differential expression. Whether the timing differences observed here translate into different pathogen restriction kinetics requires microscopic time-course analysis of pathogen growth and host cell responses on the same genotypes.

Sequencing batch was partly aligned with time point (for example, the *Rpv12* 0 hpi and 24 hpi libraries were sequenced in different batches). Batch was included in all DESeq2 models, but the ComBat-corrected values used for PCA, AED and the networks were adjusted with the condition labels as protected covariates and can overstate group separation (see Methods). Each genotype was represented by three plants that were sampled at all three time points, which limits statistical power and makes the time points of a genotype non-independent. Finally, the design lacks time-matched mock-inoculated samples, so responses to handling and time of day cannot be fully separated from pathogen-specific responses in any of the temporal changes.

## CONCLUSIONS

This transcriptome-based study shows that grapevine genotypes carrying different combinations of *Rpv* loci differ in their transcriptional profiles before and after *P. viticola* inoculation, and that part of this difference is already present at the time of inoculation. Transcriptional organization differed among the examined resistance genotypes: the triple-locus genotype changed less relative to its own 0 hpi state than the single- and double-locus genotypes. Per-genotype co-expression density covaried with the magnitude of temporal change (AED), as expected for nine-sample networks dominated by time-point means, and is therefore not independent evidence. In the *Rpv12* genotype examined here, transcripts of LRIP1/SOBIR1-related receptor genes and of EDS1– SAG101–PAD4 family members were elevated at inoculation. A structural screen of their protein interactions depended on the gene models used: receptor pairs predicted from the reference models were not reproduced with longer models from newer annotations, so verified gene models are needed before such predictions can guide experiments. Because the lines are not isogenic and no disease phenotypes (beyond breeders’ field ratings) or protein-level data were collected, these results cannot attribute effects to individual loci or establish resistance mechanisms. Rather, they show that the examined breeding genotypes have distinct transcriptional states whose underlying genetic contributions remain unresolved, and they argue for treating genotype as a primary variable, rather than a nuisance factor, when interpreting transcriptomic studies of grapevine immunity and designing pyramided cultivars.

## METHODS

### Plasmopara viticola inoculum

*P. viticola* was obtained in June 2022 as a natural inoculum from Vinselekt Michlovský a.s. vineyards in the phase of oil spots on the leaves. These leaves were stored in the dark for 48 hours at 100% humidity and 22°C until sporulation occurred. The identity of the inoculum as *Plasmopara viticola* was also confirmed by a PCR test and sequenced [73]. Excised *P. viticola* segments were incubated in water for 4 hours to prepare an inoculum, which was sprayed on sterile-washed ’Pinot Noir’ leaves. Plants were kept at 22°C and 100% humidity until sporulation, after which the densest segments were aseptically excised. Spore concentration was measured using Cellometer K2 and adjusted to 50,000 spores/ml.

### Plant material

The resistant genotypes are breeding materials developed partly within the joint grapevine breeding program of the University of Udine and the Institute of Applied Genomics (Udine, Italy) and partly by Vinselekt Michlovský a.s. (Czech Republic). All three derive from crosses involving ’Pinot Noir’, which also served as the susceptible control. The breeding line 99-1-48 was the common donor of the *Rpv1* and *Rpv12* loci, and the cultivar ’Laurot’ contributed *Rpv3*. Genotypes are designated by the resistance loci they carry (*Rpv12*, *Rpv12+1* and *Rpv12+1+3*). Further pedigree details cannot be disclosed because the cultivars are currently in a patenting process.

In the breeders’ field evaluations (scale 0 to +++), *Rpv12* is rated +++ for downy mildew and low for powdery mildew, and *Rpv12+1* is rated +++ for both. *Rpv12+1+3* has so far shown no symptoms of either disease in an unsprayed seedling population, a provisional +++ rating not yet confirmed at other sites. These ratings are breeders’ observations and were not quantified in this study.

### Plant artificial inoculation and sample collection

In January 2022, segments of one-year-old wood were collected from the vineyards of Vinselekt Michlovský a.s and planted. Genotypes with a unique combination of R-loci (*Rpv12*, *Rpv12+1*, and *Rpv12+1+3*) were selected based on laboratory analysis using SSR markers linked to resistance loci [12, 74]. ’Pinot Noir’ was used as the susceptible control and source of fresh inoculum. At the five-leaf stage, 20 µl of spore suspension was pipetted onto the abaxial side of three leaves on each plant. Leaves were then collected at three time points: 0 hours (at inoculation), 6 hours, and 24 hours post-inoculation (hpi). A one-centimeter-diameter disc was excised from the inoculation spot of each leaf and stored at -80°C until RNA isolation. Samples were collected to ensure three biological replicates for each variant.

### Total RNA isolation and sequencing

Total RNA was isolated using the Spectrum™ Plant Total RNA Kit (Sigma-Aldrich) protocol. The concentration of each sample was measured using SPECTROstar Nano. RNA integrity was measured using an Agilent 2100 Bioanalyser. The library preparation and sequencing for 36 samples, consisting of four genotypes at three time points with three replicates each, were conducted by Novogene in the UK in two batches.

### Quality control and mapping

The quality of the transcriptomic libraries was assessed using FastQC (version 0.12.1) [75] and trimmed with Trimmomatic (version 0.39) [76].

All high-quality reads were mapped onto the reference genome of *Vitis vinifera* subsp. vinifera (ENSEMBL, ID: PN40024 version 4 [37]; 35,134 annotated protein-coding genes) using the STAR mapping tool (version 2.7.4a) [77]. The featureCounts function (package Subread, version 2.0.3) [78] estimated the raw counts of mapped reads per gene. Transcripts were annotated by matching gene/protein names from multiple reference resources using BLAST’s blastP (v2.12.0+) [79], yielding a conversion table of IDs, names, and symbols.

To benchmark mapping efficiency against independently generated transcriptomes, publicly available raw RNA-seq reads from three additional studies were reprocessed through the same quality-control, trimming, alignment, and quantification pipeline used for this study’s own data (FastQC, Trimmomatic, STAR, featureCounts): two grapevine studies, Chitarrini et al. [29] (ENA study PRJEB28042; 33 samples declared, 29 successfully reprocessed) and Shi et al. [38] (SRA BioProjects PRJNA862686, PRJNA1121615, and PRJNA1118503; 83 samples declared across the three BioProjects, 74 successfully reprocessed), and one isogenic *Arabidopsis thaliana* Col-0 study, (Froussios et al. [43]; two independent experiments, ExpA and ExpB; 13 of the 14 wild-type replicates, with replicate 11 excluded as in the original study). Uniquely mapped-read proportions were compared descriptively across all four datasets (this study’s own plus the three reprocessed studies) as an independent check of data quality (Additional file 1: Table S1).

### Raw count normalization and batch correction

To verify the presence of batch effects, we compared sequencing and mapping metrics between libraries produced in two independent batches. The proportions of forward-only reads, reverse-only reads and read pairs with both mates retained after trimming differed between batches (GLM, FDR = 0.003–0.014), whereas the proportion of dropped reads did not (FDR = 0.077; Additional file 2: Figure S1).

Seven combinations of raw counts, size-factor normalization and rlog transformation, each without or with ComBat (unprotected or condition-protected), were compared by principal component analysis and the Spearman correlation between PC1 scores and sequencing batch (Additional file 3: Figure S2). Two complementary normalization strategies were then used depending on the analysis goals. For computation of aggregated expression divergence, size-factor–normalized counts from DESeq2 [39] were log₂-transformed and corrected for sequencing batch using ComBat (sva package) [41], with genotype × time point supplied as protected covariates so that differences between conditions were not removed. These values were used to compare genome-wide transcriptional distances across genotypes and time points. We used a regularized log (rlog) transformation, followed by the same condition-protected ComBat correction, for correlation analyses, clustering, and visualization. The rlog transformation stabilizes variance and improves homoscedasticity, which is desirable for assessing co-transcriptional structure but would artificially compress transcription amplitudes if used for divergence quantification. The mean–variance relationship of both corrected matrices was assessed as the Spearman correlation between gene-wise mean and variance across all 36 libraries (26,169 genes). Because ComBat was run with genotype × time point as protected covariates, downstream analyses of the corrected matrix (PCA, AED and correlation networks) can overstate group separation, particularly where the batch–condition layout is unbalanced [80]; DESeq2 inference therefore used raw counts with batch in the design.

### Analysis of expression divergence

Aggregated Expression Divergence (AED) was defined as the overall transcriptional distance between two groups of libraries. For each gene, we computed the difference between the mean batch-corrected log₂ expression of the two groups, and AED was the mean squared difference across all genes. Each resistant genotype was compared with the susceptible reference at the same time point, and each genotype was compared with itself between pairs of time points (0->6 hpi, 0->24 hpi and 6->24 hpi). AED was used as a descriptive distance. No significance tests were applied, because with three libraries per group, the same plants sampled at all time points and ComBat fitted once with the observed condition labels as protected covariates, permutation p-values would be neither informative nor calibrated.

To characterize the extent to which AED was driven by a small subset of genes versus distributed broadly across the transcriptome, we performed a gene-concentration decomposition: for each comparison, per-gene squared differences were ranked in decreasing order, and the cumulative proportion of total divergence was calculated to determine the number and percentage of genes required to account for 50%, 75%, 80%, 90%, and 95% of the total observed AED.

To show the AED and gene-concentration values expected under a specified noise model, independent Gaussian noise was added to each gene’s expression value, scaled to a fixed percentage (0.5-10%) of that gene’s own standard deviation across samples, and the same AED and gene-concentration procedures were applied to the resulting noise-injected data.

To contextualize the magnitude of the observed divergence against independent datasets, this AED framework was applied identically to three independent, publicly available transcriptomic datasets reprocessed for this purpose using the same alignment and quantification pipeline described above: an isogenic *Arabidopsis thaliana* Col-0 dataset sequenced in two independent experiments [43], a non-isogenic grapevine introgression system spanning three SRA BioProjects (Shi et al. [38]; PRJNA862686, PRJNA1121615, PRJNA1118503; Additional file 20: Table S12), and an independent *Rpv12*-carrying grapevine dataset (Chitarrini et al. [29]; ENA study PRJEB28042).

### Between-replicate variance and reference-bias assessment

We assessed whether genome-wide between-replicate variance, which includes both biological and technical variation, increased with introgression dosage. Per-gene variance across the three biological replicates was calculated within each genotype × time point combination using size-factor normalized, batch-corrected data. Mean transcriptional variance was modeled as a function of introgression dosage (0-3 major loci) and time using linear regression, with significance assessed via permutation testing (10,000 permutations). To evaluate potential reference-genome bias, the uniquely mapped-read proportion of each library was modeled with a quasibinomial generalized linear model with sequencing batch, time point and genotype as predictors, and the genotype term was assessed with an F-test.

### Differential expression analysis

Differential gene expression analysis was performed using DESeq2 [39] on raw gene counts of the genes with a total count of at least 15 across all 36 libraries (26,169 genes), with batch included as a covariate in the design formula (∼batch + condition) to account for the sequencing-batch effect identified during quality control. For each of the nine resistant-genotype x time point conditions, log2 fold changes relative to the susceptible-genotype reference at the matching time point were estimated using the apeglm shrinkage estimator [81] to stabilize fold-change estimates for low-count genes, and statistical significance was assessed using DESeq2’s Wald test against a zero-log-fold-change null, with Benjamini-Hochberg FDR correction applied across genes within each condition. A gene was considered differentially expressed in a given condition if FDR < 0.05 and |log2 fold change| > 1; the full set of 9,459 genes meeting this criterion in at least one of the nine conditions was used as the differentially expressed gene panel throughout this study. The resistant genotypes were compared directly with each other at each time point from the same fitted model, with the reference level changed for each comparison (Additional file 19: Table S11). The temporal consistency of each resistant genotype’s DE gene set was summarized as the Jaccard overlap between the sets called at each pair of time points and as the proportion of the 0 hpi set that was also called at 24 hpi (Additional file 5: Table S2).

### Comparison with an independent *Rpv12* dataset

As a quality check, the reprocessed Chitarrini samples were rlog-transformed and clustered by their pairwise Pearson correlations (Euclidean distance between correlation profiles, complete linkage; Additional file 14: Figure S6). For the cross-study comparison, inoculated samples of Chitarrini et al. [29] at 12 h and 24 h were each compared with mock samples at 0 h (DESeq2, ∼condition, three replicates per group; same apeglm shrinkage and DE criterion). The corresponding *Rpv12* contrasts of this study (6 vs 0 hpi and 24 vs 0 hpi) were extracted from the full model above (∼batch + condition, all 36 libraries), because the *Rpv12* 0 hpi and 24 hpi libraries were sequenced in different batches. A sensitivity model additionally included the individual plant nested within genotype (∼batch + genotype + genotype:plant + genotype:time), because the same three plants per genotype were sampled at all time points. Within the 24,338 genes tested in both studies, the overlap between DE gene sets was tested with a one-sided hypergeometric test and the directional agreement of shared genes with a one-sided Fisher’s exact test of the 2 × 2 table of up-and downregulation calls in the two studies, which keeps each study’s proportions of up-and downregulated genes; the agreement expected by chance was calculated from these proportions. GO biological-process enrichment of the direction-concordant genes was tested with topGO [82] (Fisher’s exact test, Benjamini–Hochberg adjustment), using the TAIR GO annotation of each gene’s closest *Arabidopsis* homolog and the genes tested in both studies as background.

### Gene categorization

We categorized each differentially expressed gene (DEG) in every resistant genotype according to its specific time-dependent transcriptional pattern. At each time point (0, 6, and 24 hpi), the gene is classified as not differentially expressed relative to the reference (E or 0; the DE criterion was not met, which does not demonstrate equal expression), downregulated (D or -1), or upregulated (U or 1). The transcriptional dynamics of each gene in a particular genotype are represented by a combination of three characters (E or 0, D or -1, U or 1). These three character combinations of each genotype (the simplified transcript abundance pattern) allow us to categorize the dynamics of transcriptional changes into six categories: 1) changes detected at 0 hpi, or at 0 and 6 hpi, are labeled Initial Expression Variation (IEV); 2) changes detected at 6 and 24 hpi, Early Response (ER); 3) changes detected only at 6 hpi, Transient Stress Response (TSR); 4) changes detected only at 24 hpi, Late Response (LR); 5) changes detected at all three time points, Sustained Change (SCh); and 6) all other patterns, Complex Pattern (CP). All DE calls within categories 1–5 must have the same direction (for example, UUU or DDD for SCh); patterns that switch direction (for example, UDE or UUD) are classed as CP. These category names are operational labels for patterns of DE calls; they do not by themselves demonstrate a physiological response.

### Grouping of gene categories

Categorized genes can be further grouped by comparing the simplified transcript abundance pattern among genotypes. A pattern is considered shared only when the three-letter pattern is identical in all genotypes with DE calls; if a pattern is shared between two genotypes, the third genotype has no DE call at any time point (EEE), and all other combinations form Group VI. We define six groups (Additional file 11: Figure S5): Group I -pattern shared by all three genotypes, Group II -pattern shared by *Rpv12* and *Rpv12+1* genotypes, Group III -pattern shared by *Rpv12+1* and *Rpv12+1+3* genotypes, Group IV -pattern shared by *Rpv12* and *Rpv12+1+3* genotypes, Group V -genotype-specific gene categories (changes found in only one genotype), Group VI -complex patterns across genotypes. Group V has three subgroups: (a) *Rpv12*-specific changes, (b) *Rpv12+1*-specific changes, and (c) *Rpv12+1+3*-specific changes. Because each call is made against the susceptible reference, a gene that is DE in one genotype but not in another has not been tested for a difference between those two genotypes; group assignments describe patterns of calls, not tested contrasts.

### Statistical models of DEG counts

To assess how genotype, time point and direction of regulation influence the number of DEGs, we applied generalized linear models (GLMs). For DEG counts, we used the negative binomial GLM with a log link. Because DEG counts were derived from normalized transcript abundance estimates relative to the susceptible genotype, an additional library-size offset was not included in the generalized linear models. For Models E and F, the direction of regulation of each gene was taken from its first DE call or, for complex patterns (CP), from the majority of up and down calls (ties counted as up). Models E and F include the genes of Groups I–V (5,238 of 9,459 differentially expressed genes); Group VI (4,221 genes, 44.6%) has no shared or genotype-specific pattern and is excluded.

Count models (A–E) used the negative binomial family; Model F, which models proportions of each group’s gene pool, used a quasibinomial family. To assess whether DEG counts depended on single libraries, the DESeq2 model was refitted 36 times, each time omitting one library, with the same design and differential-expression criterion, and DEG counts per genotype and time point were compared across refits.

### Pearson correlation network analysis

Co-transcriptional modules were defined for each gene of interest as that gene together with every other gene, among the full 26,169-gene expression matrix, positively correlated with it at r ≥ 0.8077 (Pearson correlation, rlog-transformed and condition-protected-ComBat-corrected values), provided at least three such correlated partners were found (four genes total, including the gene of interest). This threshold corresponds to the top 0.5th percentile of the full distribution of 342,395,196 unique pairwise correlations among all 26,169 detected genes, re-derived directly on the condition-protected ComBat matrix used throughout this study. Modules were built for the full set of 9,459 canonical-DESeq2 differentially expressed genes (above).

The robustness of this network definition to methodological choices was assessed in several complementary ways. The network was rebuilt at the top 1%, 0.5% and 0.1% of pairwise correlations (r ≥ 0.757, 0.8077 and 0.8839), with all 9,459 differentially expressed genes as anchors at each threshold. At each threshold, network edges (anchor–partner pairs) were compared with STRING-db v12.0 links for *Vitis vinifera* [83] under three evidence definitions, each at score ≥ 400: the combined score; the experimental channel only; and the combined score recomputed without the text-mining and curated-database channels (STRING’s channel-combination scheme with a prior of 0.041). The fraction of supported edges was compared with that of 2 × 10⁶ random pairs of a differentially expressed gene and any STRING-mapped gene (fold enrichment). The stability of correlation-based edges under simulated measurement noise was assessed on the primary network matrix (rlog-transformed, condition-protected ComBat-corrected values of the 9,459 differentially expressed genes).

Independent Gaussian noise with a standard deviation of 0.5%, 1%, 2%, or 5% of each gene’s own standard deviation across the 36 samples was added (200 replicates per level), and the network of all gene pairs was rebuilt at r ≥ 0.8077 after each perturbation. Stability was quantified as edge recall (the fraction of edges in the noise-free network that were retained) and as the Jaccard similarity between the perturbed and noise-free edge sets. The same procedure applied to DESeq2 negative-binomial Pearson residuals (design ∼ batch + condition) of the same genes served as a separate sensitivity analysis on a different expression representation.

Module gene sets were compared with STRING-db (v12.0) [83] interaction and co-expression evidence; for the modules at r ≥ 0.8077 and for genome-wide WGCNA modules, the proportion of modules with elevated STRING scores (empirical p < 0.05 per module, size-matched permutation test with 200 random gene sets per size class) is reported descriptively, because overlapping modules are not independent.

Classical weighted gene co-expression network analysis (WGCNA) was performed on the genome-wide condition-protected ComBat matrix following Langfelder and Horvath [84], with a signed network, soft-thresholding power β = 8 (selected with pickSoftThreshold() using the approximate scale-free topology fit criterion, R² ≈ 0.85) and topological-overlap-based module detection, as a comparator for module size and within-module coherence (PC1 variance).

### Analysis of network structure

To examine how transcriptome-wide network architecture changes over time and with increasing major resistance locus dosage, we constructed correlation-based co-transcriptional networks (igraph R package [85]) separately per genotype and per time point using the 9,459 DE genes. For each subset of samples (e.g. *Rpv12*, *Rpv12+1, Rpv12+1+3*, or susceptible; and separately 0 hpi, 6 hpi, 24 hpi), we computed pairwise Pearson correlations on rlog-transformed, condition-protected ComBat-corrected expression values. An edge was retained if the correlation exceeded a stringent threshold (r ≥ 0.8077; 99.5th percentile of the global correlation distribution; networks built at r ≥ 0.757 and r ≥ 0.8839 gave the same order of genotypes and of time points; Additional file 16: Table S9). Logical adjacency matrices were converted into undirected igraph objects with edge weights equal to the corresponding correlation coefficients. Global network metrics were then extracted using igraph: number of nodes, number of edges, density (edge_density), mean degree, and clustering coefficient (transitivity(type="global")). Network density was regressed against the number of major resistance loci and evaluated with Pearson and Spearman correlations, and was also regressed against time point. Per-genotype networks (nine samples) and per-time-point networks (twelve samples) differ in sample size, so a fixed r threshold corresponds to different null exceedance rates (t = r√(n−2)/√(1−r²)); densities were therefore compared only within each network type.

## Functional enrichment analysis

Functional enrichment of Gene Ontology (GO) terms, Reactome pathways, UniProt keywords, and protein domains was assessed using the STRING database version 12.0 [83], with *Vitis vinifera* annotation as well as *Arabidopsis* homologs. Principal component analysis used the rlog-transformed, condition-protected ComBat-corrected values of all 26,169 genes; for PC1–PC4, the 50 genes with the highest contribution (squared loading, normalized per component) were tested with their *Arabidopsis* homologs (Additional file 8: Table S4). The 18 immunity-related genes of Figure 5 were chosen from genes with literature-established immune functions [47–52] that differed between *Rpv12* and the susceptible control at 0 hpi, together with the genes of the predicted networks; their *Arabidopsis* homologs were queried in STRING (combined score ≥ 0.4).

### Protein–protein interaction prediction

To screen for candidate protein interactions, we performed systematic AlphaFold2-Multimer [86, 87] structural predictions using ColabFold [88]. Candidate pairs came from three sources: a gene of interest and members of its co-transcriptional module (for some genes of interest, as defined in an earlier version of the network built on the unprotected ComBat matrix); pairs chosen from the literature or from shared transcriptional patterns, including a screen of all pairwise combinations of BRI1, BAK1, CLV1 and BAM1 homologs and leucine-rich repeat genes of the SOBIR1 and LRIP1 modules (receptor-family screen); and cross-module pairs among immunity-related proteins from different modules. The source of each pair is given in Additional file 18: Table S10; all pairs were scored with the same criteria. Protein sequences were taken from the PN40024 v4 reference annotation, so the models do not show that the proteins encoded in the examined genotypes interact. Paired FASTA sequences were generated for each combination and processed using colabfold_batch (--model-type alphafold2_multimer_v3 --num-models 5 --num-recycle 3). Each pair was evaluated across the five models produced by the five AlphaFold2-Multimer parameter sets from a shared MSA (ColabFold v1.6.2, with the underlying AlphaFold2-Multimer model version, v3). Interface quality was assessed using ipTM (interface predicted TM-score), pLDDT (per-residue confidence), interface contact count (Cα–Cα distance < 8 Å), and contact-filtered PAE (predicted aligned error, calculated as the mean PAE across residue pairs forming interface contacts). A pair was classified as a high-confidence interaction if ipTM ≥ 0.75 in all five models, with ≥ 5 interface contacts and a mean contact-filtered PAE < 10 Å across the five models; pairs meeting the single-model version of this criterion (ipTM ≥ 0.7, pLDDT ≥ 50, ≥ 5 contacts, and either contact-filtered PAE < 10 Å or ≥ 50% of interface contacts with PAE < 10 Å) in at least one of the five models but not the full five-model criterion were retained separately as partial evidence. For the SOBIR1–LRIP1 pair, interface residues (Cα–Cα distance < 8 Å, pLDDT ≥ 50 in both chains) were mapped to SOBIR1 domains (signal peptide, LRR ectodomain, transmembrane segment, juxtamembrane region and kinase domain). Because PN40024 v4 gene models are computational predictions [36], the length of each protein in the structural interaction set was compared with that of its closest *Arabidopsis* homolog (best BLASTP alignment), and models lacking more than 25 residues of the homolog at either end were flagged (24 of 43 genes; Figure 4A, C).

The behavior of the screen was checked with a pre-specified negative-control set defined before screening: four transcription-factor hubs (PRE6, WRKY51, WRKY55, NAC29) and their co-expression partners. Established literature-or homology-supported complexes (the EDS1-SAG101-PAD4 family) were screened under the same procedure and reported separately as positive controls rather than pooled with the other candidate interactions. Both control sets are computational checks; they are not experimentally verified negatives or positives and cannot establish experimental specificity. Six annotated EDS1-family paralogs at this locus were included in the screen; Vitvi17g04216 was carried forward as the primary EDS1 node based on its upregulation in *Rpv12* at 0 hpi. Cross-module pairings between hub proteins from different co-transcriptional modules were also screened and are reported descriptively as part of the discovery screen.

To assess whether predicted interactions reflect generic AlphaFold2-Multimer "stickiness", a potential modeling artefact in which a protein forms plausible interfaces with many unrelated partners, a dedicated promiscuity panel was constructed: 100 candidate proteins (80 receptor-kinase/receptor-like-family genes, identified by keyword-matching *Arabidopsis* homolog annotations for receptor-kinase/RLK/LRR-RK/SERK/WAK/lectin-kinase-type terms, plus 20 non-receptor background genes) were each screened against 16 query proteins from the SOBIR1/LRIP1 network under the same five-model procedure and criteria described above (1,600 pairwise predictions total). A query protein was flagged as promiscuous, and excluded from consideration as a specific interaction irrespective of its raw confidence score, if it met the high-confidence criterion against multiple, functionally unrelated panel candidates. A predicted interaction was reported as a candidate interaction outside known complexes only if it (i) passed the strict five-model criterion, (ii) did not belong to an established literature-or homology-supported complex, and (iii) involved no partner flagged as promiscuous in this panel.

### Gene models and repeated structure predictions

The models of the 43 genes of the structural interaction set, and in detail those of the nine genes of the predicted LRIP1/SOBIR1 network, were compared with those of the same loci in the GenBank and RefSeq annotations of the telomere-to-telomere PN40024 assembly [89] and in Phytozome v2.1 [90]. Each v4 gene region was placed on the telomere-to-telomere assembly with minimap2 [91], which gave the sequence identity between assemblies; protein models were compared by BLASTP, and a model was counted as complete when it covered its *Arabidopsis* homolog to within 25 residues at both ends and was at most 1.25 times its length; signal peptides were predicted with SignalP 6.0 [92] (Additional file 18: Table S10, sheets gene_models_network and gene_models_loci). Introns of the longer models were compared with splice junctions in two RNA-seq libraries of this study (*Rpv12*, 0 and 6 hpi) mapped to the telomere-to-telomere assembly with STAR, distinguishing junctions supported by uniquely mapped reads from those seen only in multi-mapped reads. All ten candidate pairs were then predicted again with the longest available model of each protein (ColabFold v1.6.3, otherwise the same settings) and scored with the same criteria and script as the screen (sheet AF2_rerun_gene_models).

### Domain architecture and helper-NLR classification

Pfam-A domains were annotated for all PN40024 v4 proteins with HMMER v3.3.2 hmmscan [93] against the Pfam-A database v35.0 [94] (-E 0.001 --domE 0.001), and the primary isoform (P001) of each of the 43 genes in the structural interaction set was analyzed; hits were retained only if both the full-sequence and the independent-domain E-value were below 0.001. Overlapping hits within a protein were resolved by keeping the hit with the highest domain score, giving a non-overlapping domain set for each protein; each Pfam model was counted once per protein, and results are summarized as numbers of genes per domain category.

Helper NLRs were identified as proteins carrying the RPW8 domain (Pfam PF05659) in the genome-wide hmmscan annotation (primary isoform). Those that also carry an NB-ARC domain were classed as NRG1 or ADR1 homologs according to their best *Arabidopsis* BLASTP hit (NRG1, AT5G66900 or AT5G66910; ADR1, AT1G33560, AT4G33300 or AT5G04720); differential expression was taken from the DESeq2 analysis described above (Additional file 9: Table S5).

### Interactive database and visualization tool

To facilitate exploration of the dataset, we developed an interactive web application using the R Shiny framework [95] (shiny, ggplot2, DT, Biostrings, shinyWidgets packages). The app allows users to query gene-level information, visualize abundance profiles, inspect co-transcriptional modules, and download associated data. It integrates normalized transcription matrices (rlog-and ComBat-corrected), raw read counts, and reference FASTA sequences of *Vitis vinifera* (PN40024.v4). Homology relationships with *Arabidopsis thaliana* were obtained by BLASTP (e-value < 1 × 10⁻⁶) against TAIR10 proteins and are provided unfiltered to retain all potential homologs. Each gene entry links to UniProt, NCBI, and TAIR10 identifiers, and displays its co-transcriptional partners (Pearson’s r ≥ 0.8077). The application, titled *Grapevine Guardians: Unraveling Rpv Pyramidization’s Impact on Immunity*, is available at https://vierakovacova.shinyapps.io/playing_with_immunity/ and the complete source code is accessible via GitHub (https://github.com/vierocka/plant_immunity_Vitis).

## List of abbreviations

AED: Aggregated expression divergence
BiFC: Bimolecular fluorescence complementation
BLASTP: Protein Basic Local Alignment Search Tool
CC: Coiled-coil
CNL: Coiled-coil nucleotide-binding leucine-rich repeat receptor
CP: Complex pattern
DE: Differentially expressed / differential expression
DEG: Differentially expressed gene
EIX: Ethylene-inducing xylanase
ENA: European Nucleotide Archive
ER: Early response
ETI: Effector-triggered immunity
FDR: False discovery rate
GLM: Generalized linear model
GO: Gene Ontology
hpi: Hours post-inoculation
IEV: Initial expression variation
iPAE: Interface predicted aligned error
ipTM: Interface predicted template modelling score
log₂FC: log₂ fold change
LR: Late response
LRR: Leucine-rich repeat
MAPK: Mitogen-activated protein kinase
MSA: Multiple sequence alignment
NB: Nucleotide-binding
NB-ARC: Nucleotide-binding adaptor shared by APAF-1, R proteins and CED-4
NCBI: National Center for Biotechnology Information
NLR: Nucleotide-binding leucine-rich repeat receptor
PAE: Predicted aligned error
PAMP: Pathogen-associated molecular pattern
PC / PCA: Principal component / principal component analysis
PCD: Programmed cell death
PCR: Polymerase chain reaction
pLDDT: Predicted local distance difference test
PPI: Protein–protein interaction
PRR: Pattern recognition receptor
PTI: Pattern-triggered immunity
pTM: Predicted template modelling score
RIN: RNA integrity number
RLK: Receptor-like kinase rlog Regularized logarithm
RLP: Receptor-like protein
RNL: RPW8-type nucleotide-binding leucine-rich repeat receptor
ROS: Reactive oxygen species
*Rpv*: Resistance to *Plasmopara viticola*
SCh: Sustained change
SRA: Sequence Read Archive
SSR: Simple sequence repeat
STRING: Search Tool for the Retrieval of Interacting Genes/Proteins
TAIR: The *Arabidopsis* Information Resource
TIR: Toll/interleukin-1 receptor
TM: Transmembrane segment
TNL: Toll/interleukin-1 receptor nucleotide-binding leucine-rich repeat receptor
TSR: Transient stress response
WAK: Wall-associated kinase
WGCNA: Weighted gene co-expression network analysis

**Gene names**

SOBIR1: SUPPRESSOR OF BIR1-1
LRIP1: LEUCINE-RICH REPEAT IMMUNE PROTEIN 1
EDS1: ENHANCED DISEASE SUSCEPTIBILITY 1
PAD4: PHYTOALEXIN DEFICIENT 4
SAG101: SENESCENCE-ASSOCIATED GENE 101
BAK1: BRI1-ASSOCIATED RECEPTOR KINASE 1
BIR1: BAK1-INTERACTING RECEPTOR-LIKE KINASE 1
SERK: SOMATIC EMBRYOGENESIS RECEPTOR KINASE
NRG1 / ADR1: N REQUIREMENT GENE 1 / ACTIVATED DISEASE RESISTANCE 1
RPW8: RESISTANCE TO POWDERY MILDEW 8
ELR: ELICITIN RESPONSE
RUN1: RESISTANCE TO UNCINULA NECATOR 1
ACD6: ACCELERATED CELL DEATH 6
BAM1: BARELY ANY MERISTEM 1
BRI1: BRASSINOSTEROID INSENSITIVE 1
CLV1: CLAVATA 1
FER: FERONIA
MIK2: MDIS1-INTERACTING RECEPTOR-LIKE KINASE 2
LAZ5: LAZARUS 5
SARD4: SAR-DEFICIENT 4
GSO2: GASSHO 2
RGI1: RGF1 INSENSITIVE 1
MRH1: MORPHOGENESIS OF ROOT HAIR 1
MYB: MYB (myeloblastosis-related) transcription-factor family
NAC: NAM, ATAF1/2 and CUC2 transcription-factor family
PRE6: PACLOBUTRAZOL RESISTANCE 6
WRKY: WRKY transcription-factor family

## Declarations

### Ethics approval and consent to participate

Not applicable. The grapevine genotypes are breeding materials that were provided for this study by Vinselekt Michlovský a.s. (Czech Republic); no wild plants were collected. All experimental work with plants complied with relevant institutional, national and international guidelines and legislation.

### Consent for publication

Not applicable

### Clinical trial number

Not applicable.

### Availability of data and materials

Raw RNA-seq data have been deposited in the NCBI Sequence Read Archive under accession number NCBI BioProject: PRJNA1358055. Processed data — including DESeq2 differential expression results, AlphaFold2-Multimer-tested PPIs, raw counts, ComBat batch-effect-corrected counts, and sample metadata — have been deposited in Zenodo under DOI 10.5281/zenodo.21623285. All supplementary tables (Tables S1– S12), including the reprocessed independent RNA-seq datasets used for cross-study comparison [29, 38, 43] (per-sample DESeq2 size factors, normalized expression values and differential expression results; Additional file 6: Table S3; Additional file 13: Table S8; Additional file 20: Table S12), have been deposited in Zenodo under DOI 10.5281/zenodo.23579867; because of their size, Additional files 13, 19 and 20 are available only from this record.

An interactive Shiny application that mirrors and facilitates exploration of the transcriptomic results is available at https://vierakovacova.shinyapps.io/playing_with_immunity/. All scripts used for quality control, mapping, differential expression analysis, AED, and network analysis are available at https://github.com/vierocka/plant_immunity_Vitis.

### Authors’ contributions

VK: conceptualisation of the sequencing data analysis and interpretation, data curation, bioinformatic analysis, and co-writing of the original manuscript; MH: experimental work and co-writing of the original manuscript; MB: conceptualisation of the experiment and co-writing of the original manuscript; KB: conceptualisation of the experiment and experimental work. All authors read and approved the final manuscript.

## Acknowledgements

We would like to express our gratitude to VINSELEKT MICHLOVSKÝ a.s. for providing the grapevine breeding materials used in this study.

## ADDITIONAL FILES

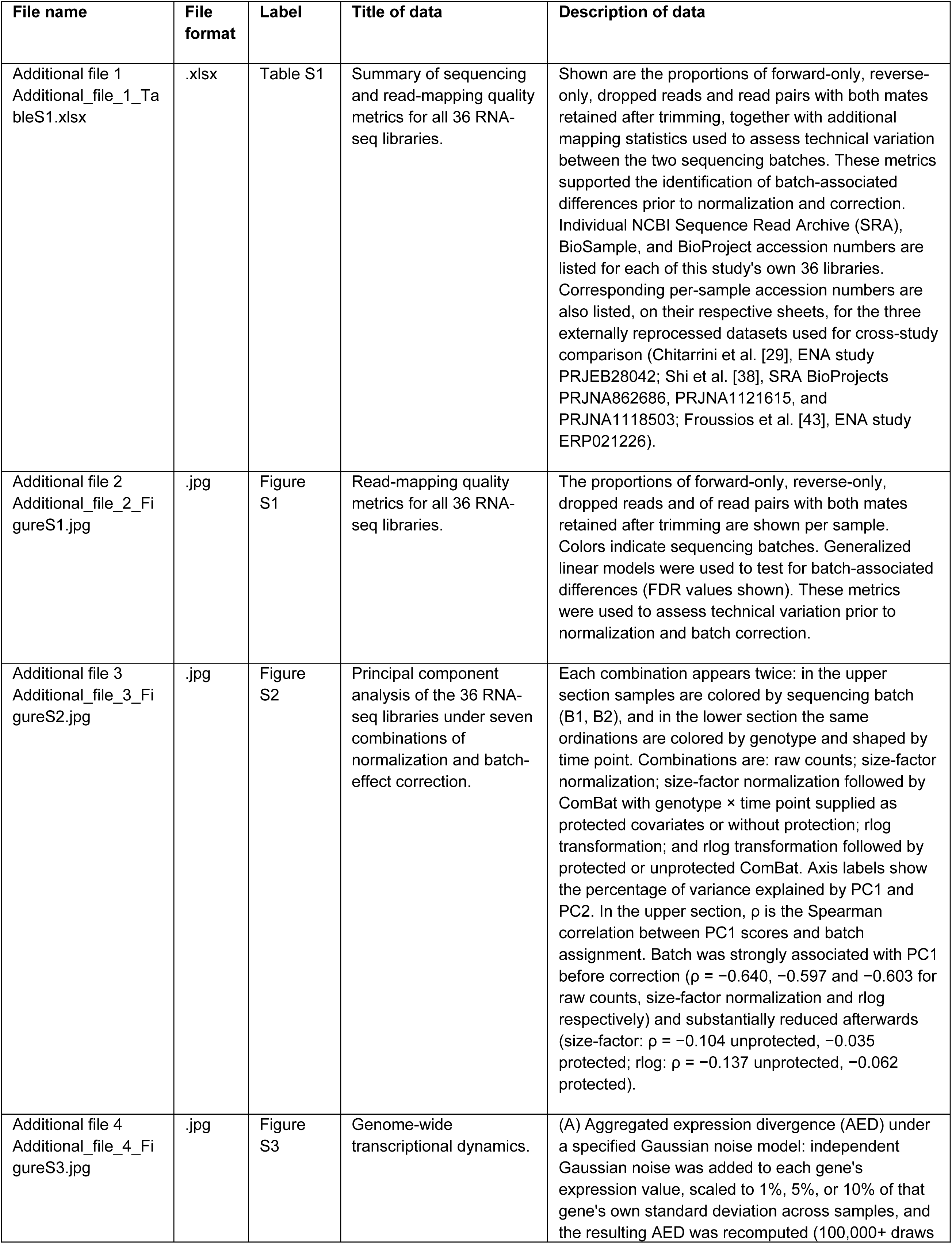
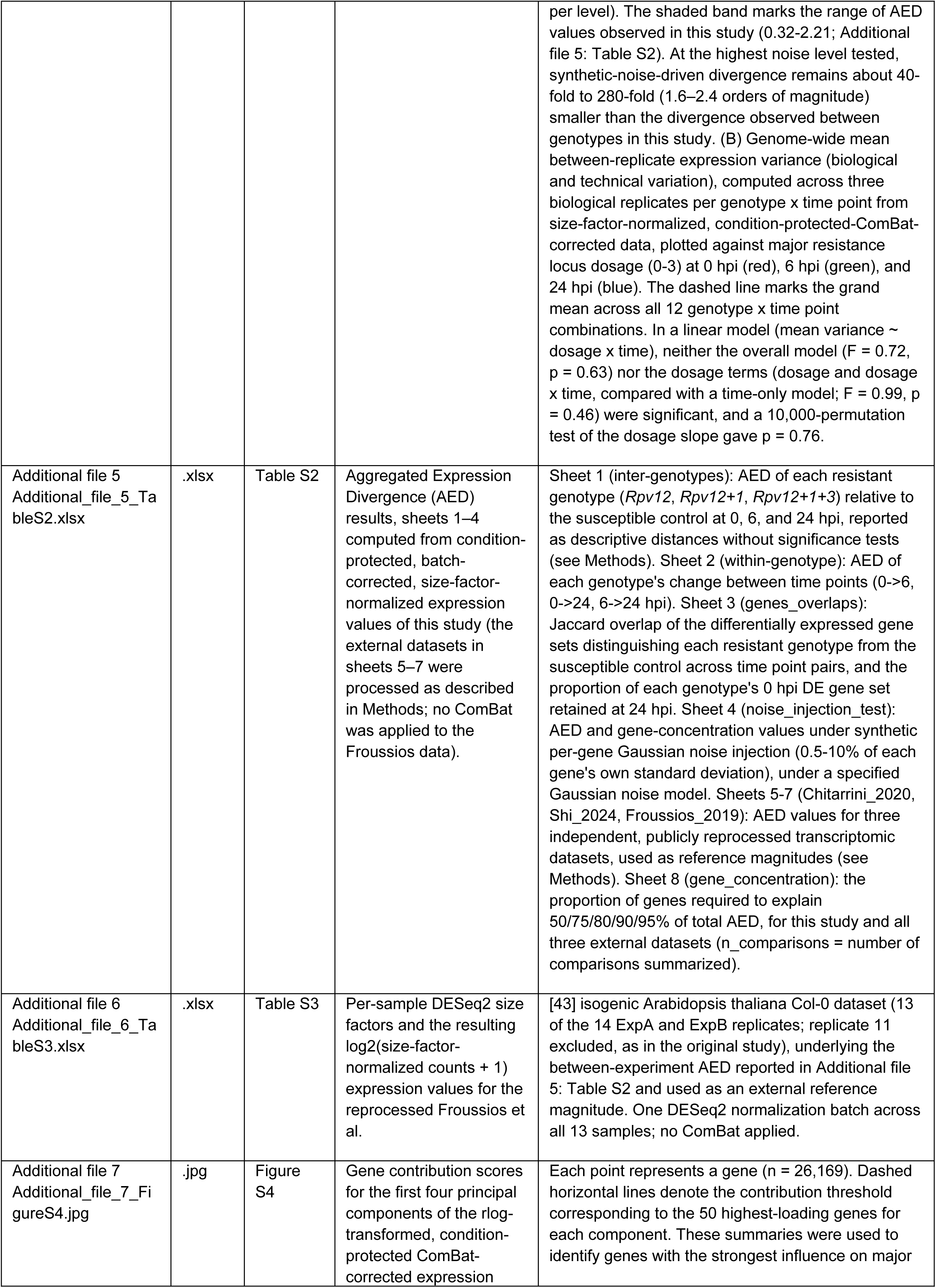
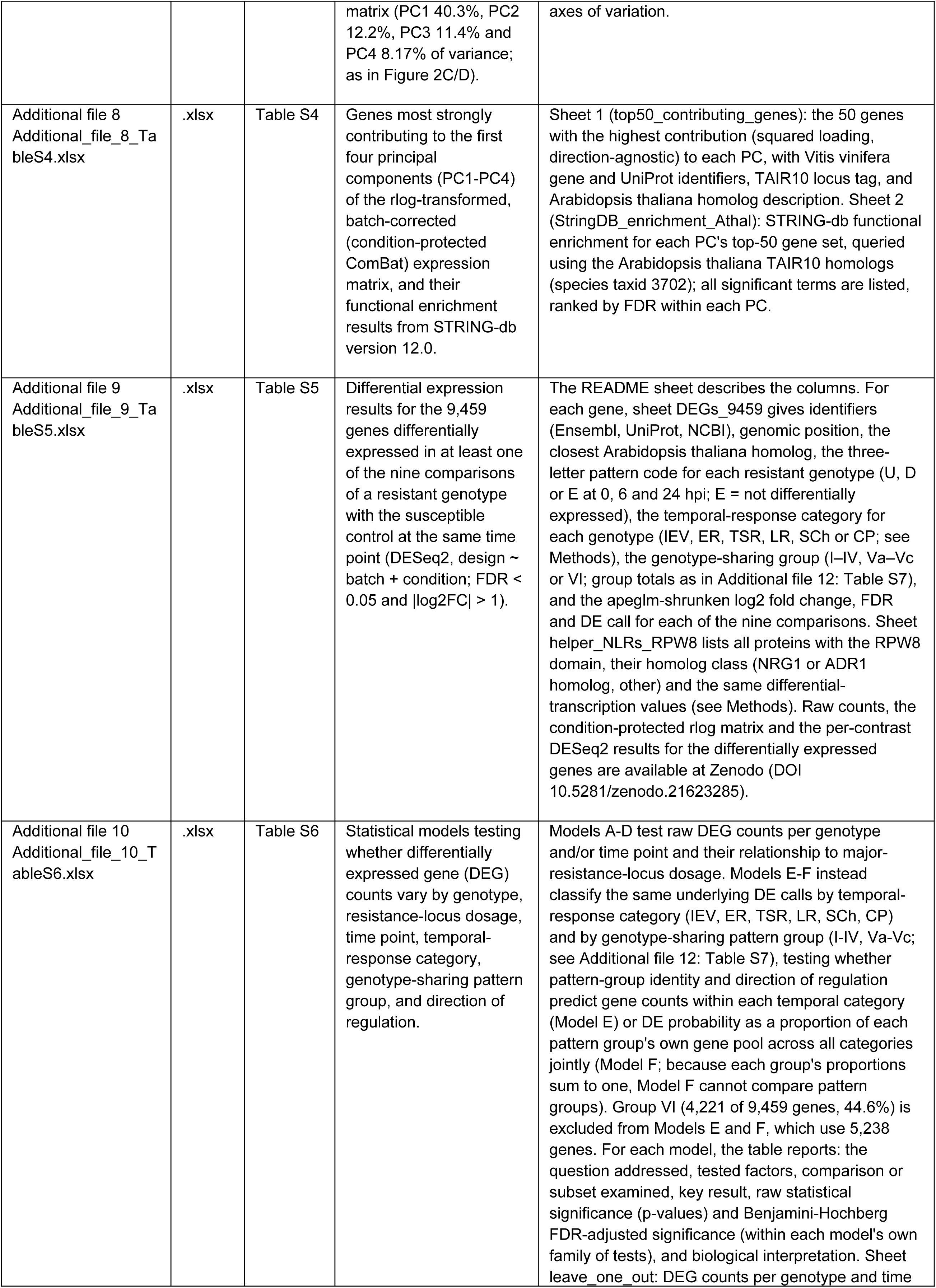
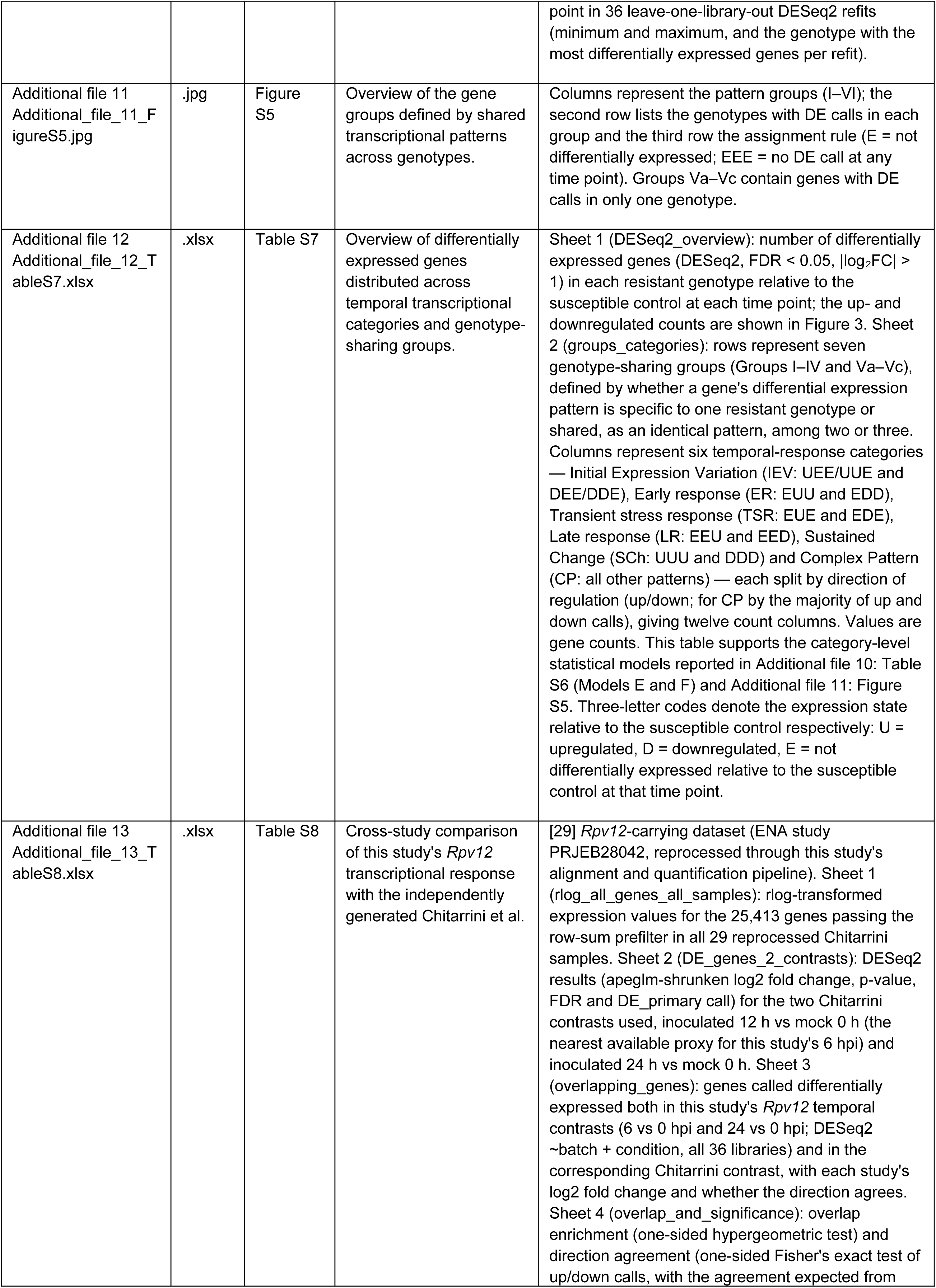
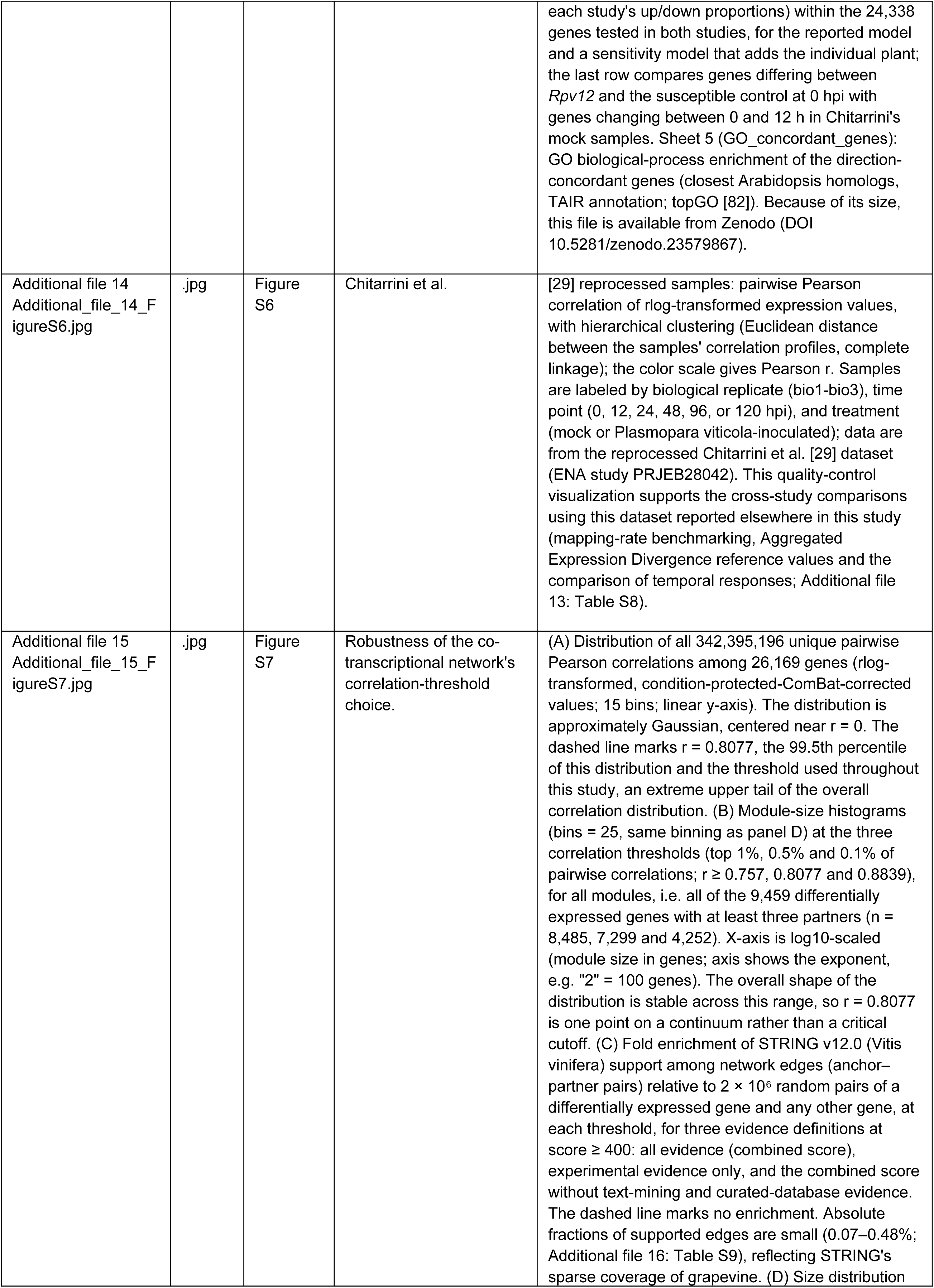
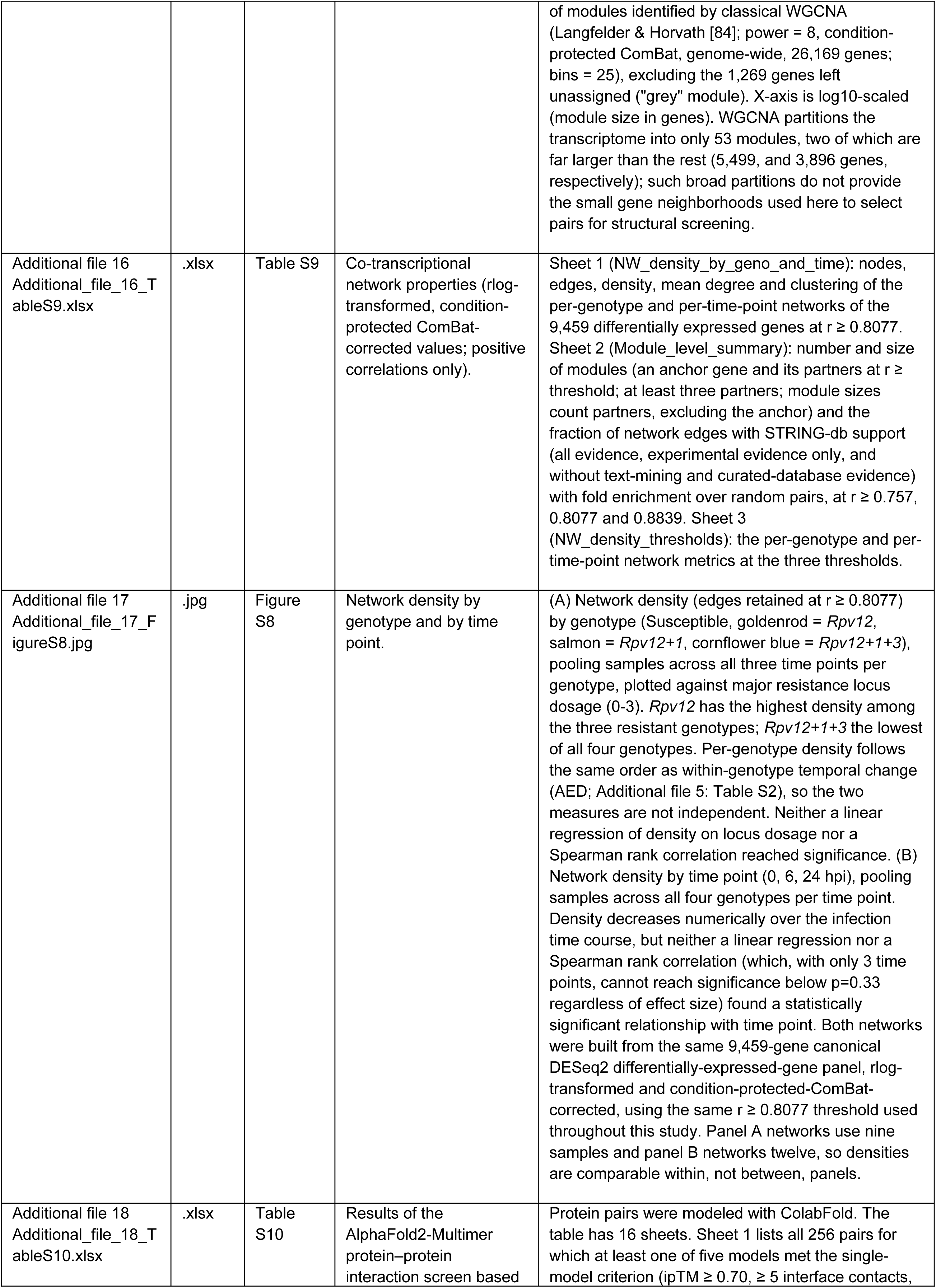
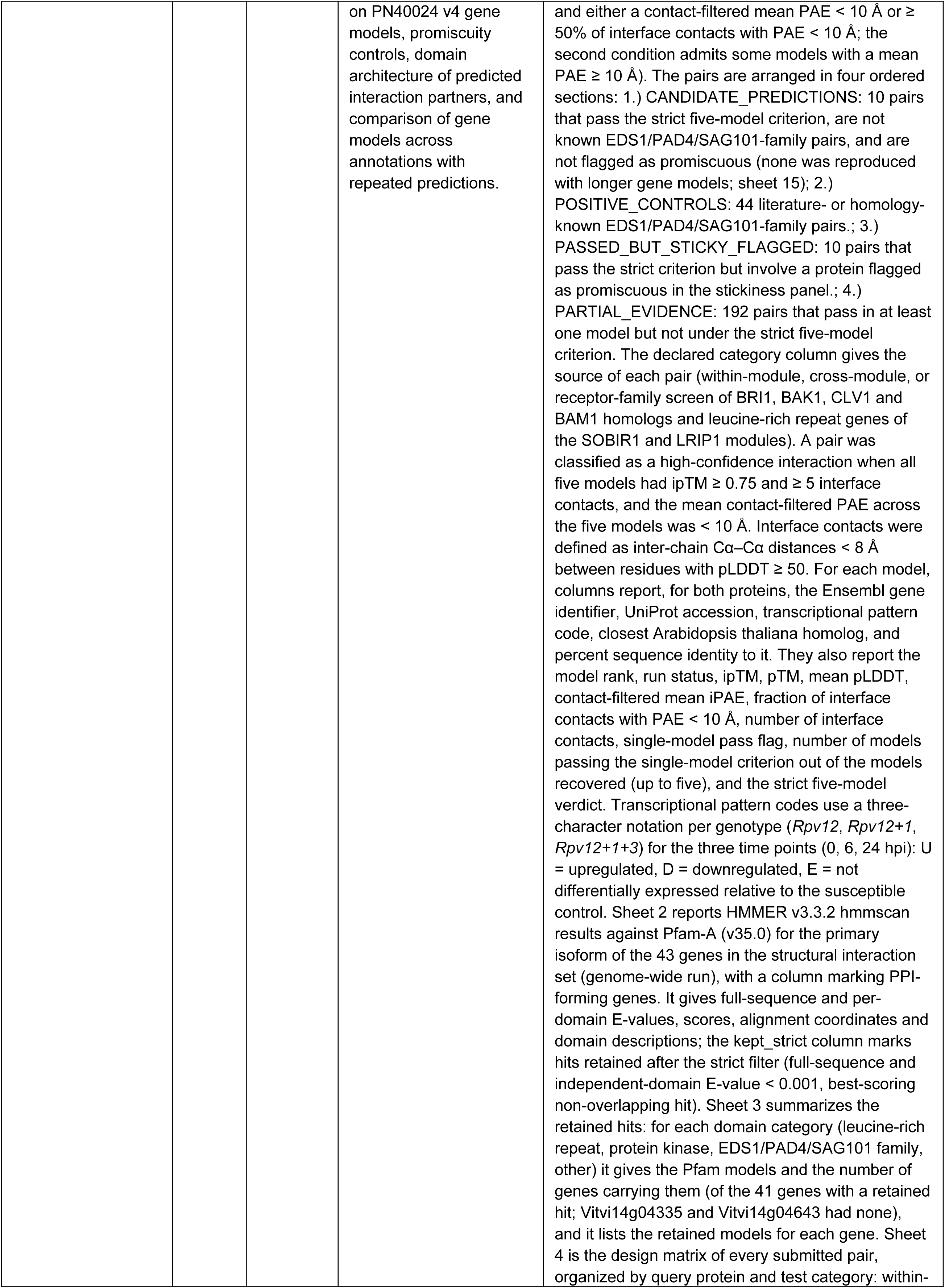
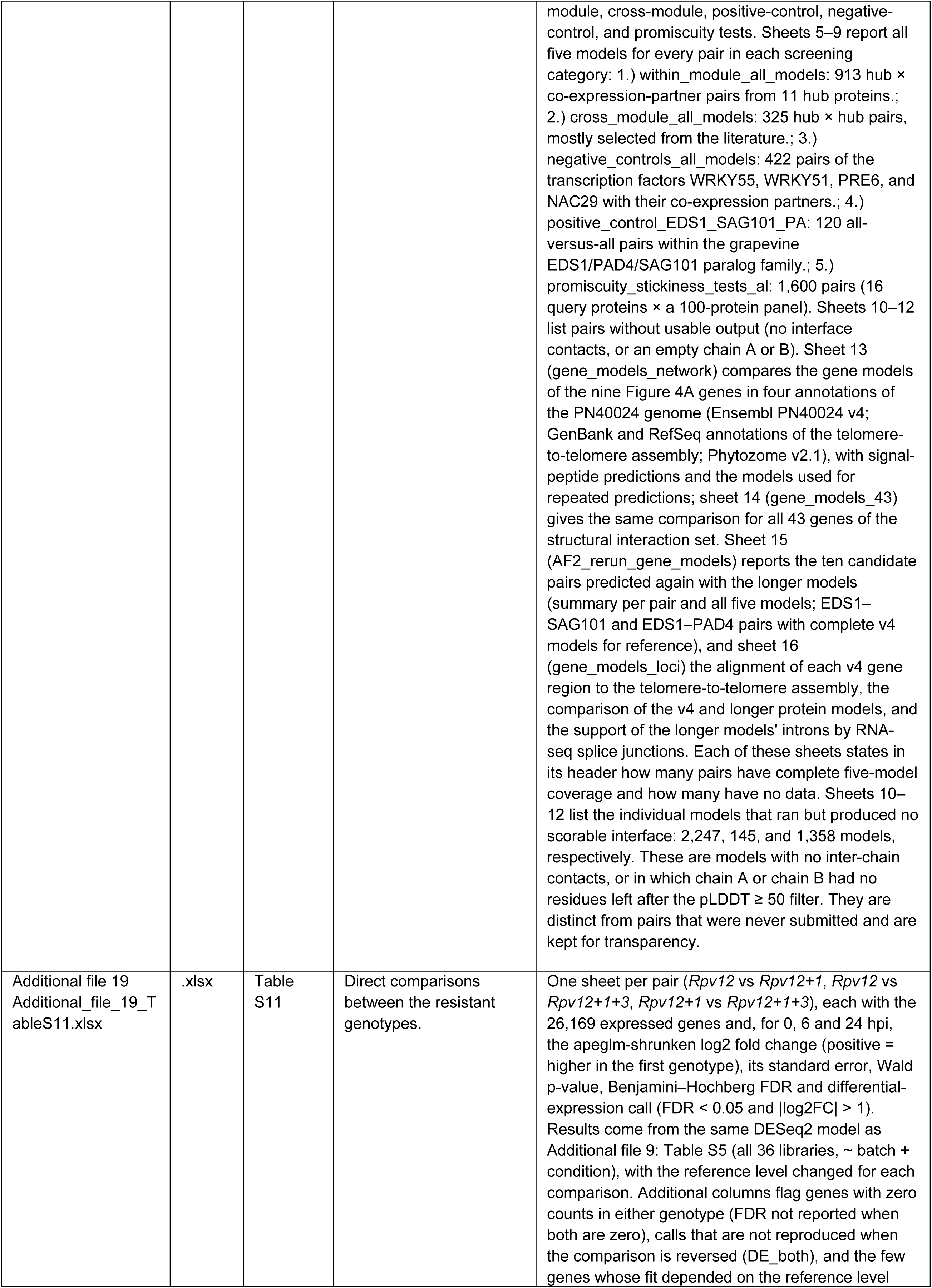
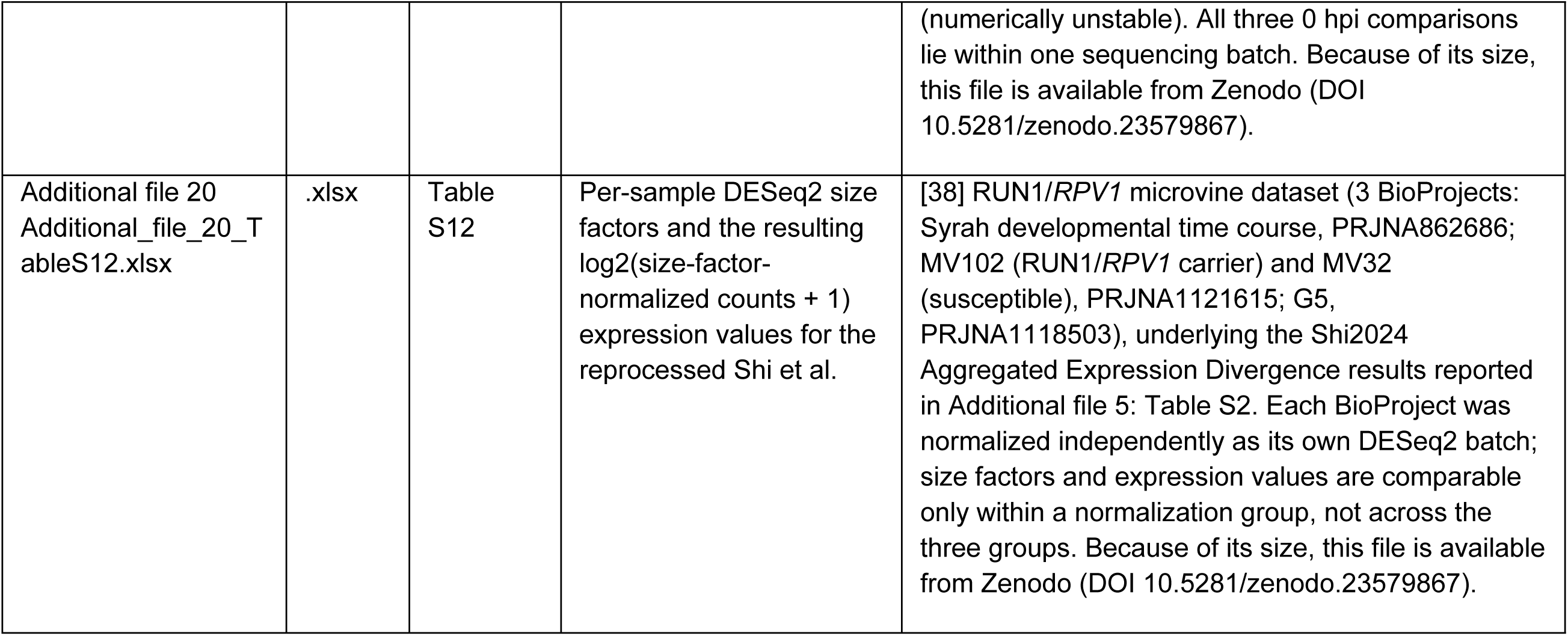

