## Supplementary figures and images for "Genotype-associated transcriptional differences among grapevine lines carrying single and pyramided *Rpv* resistance loci"

### Supplementary Figure 1

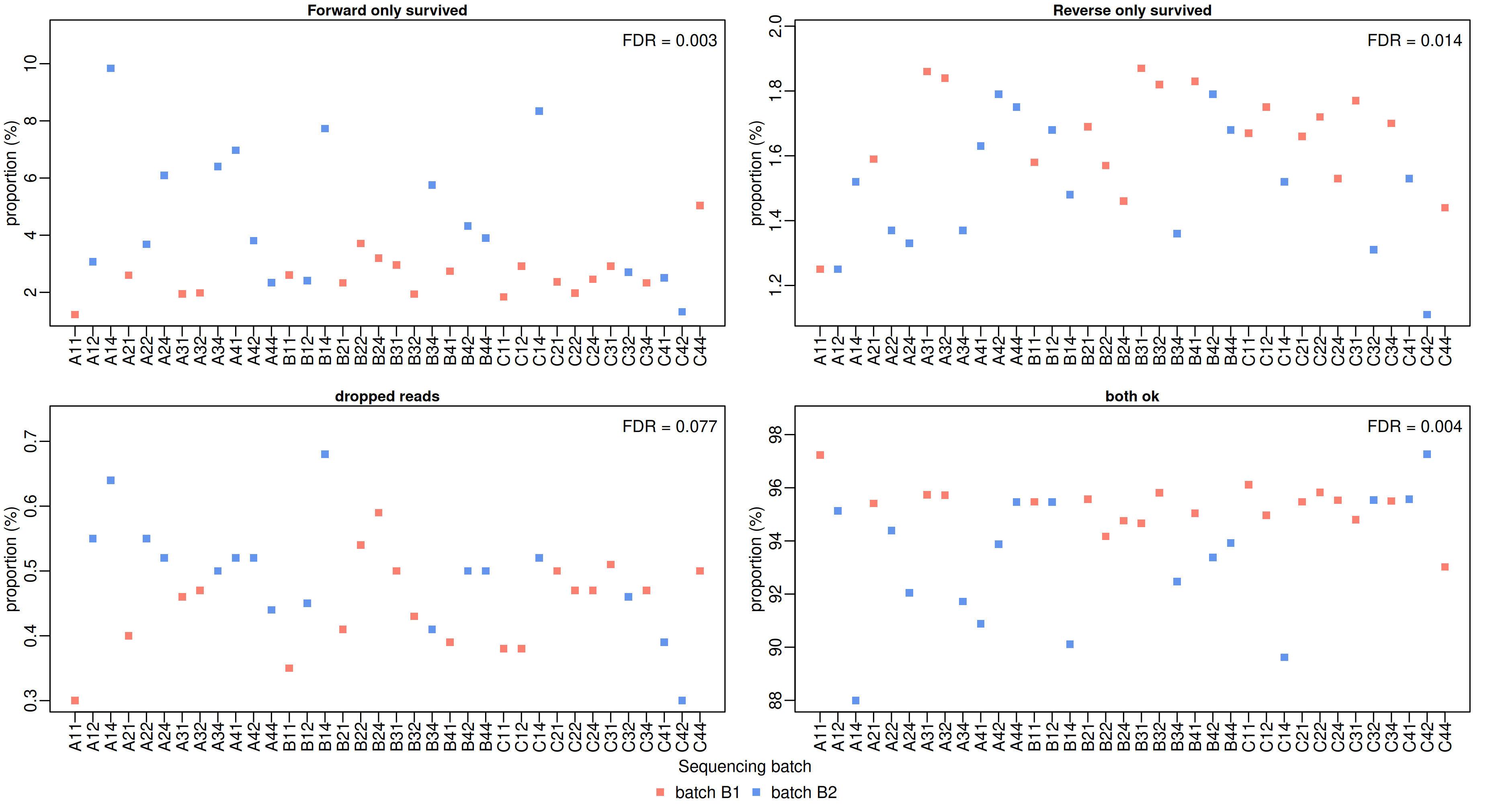

### Supplementary Figure 2

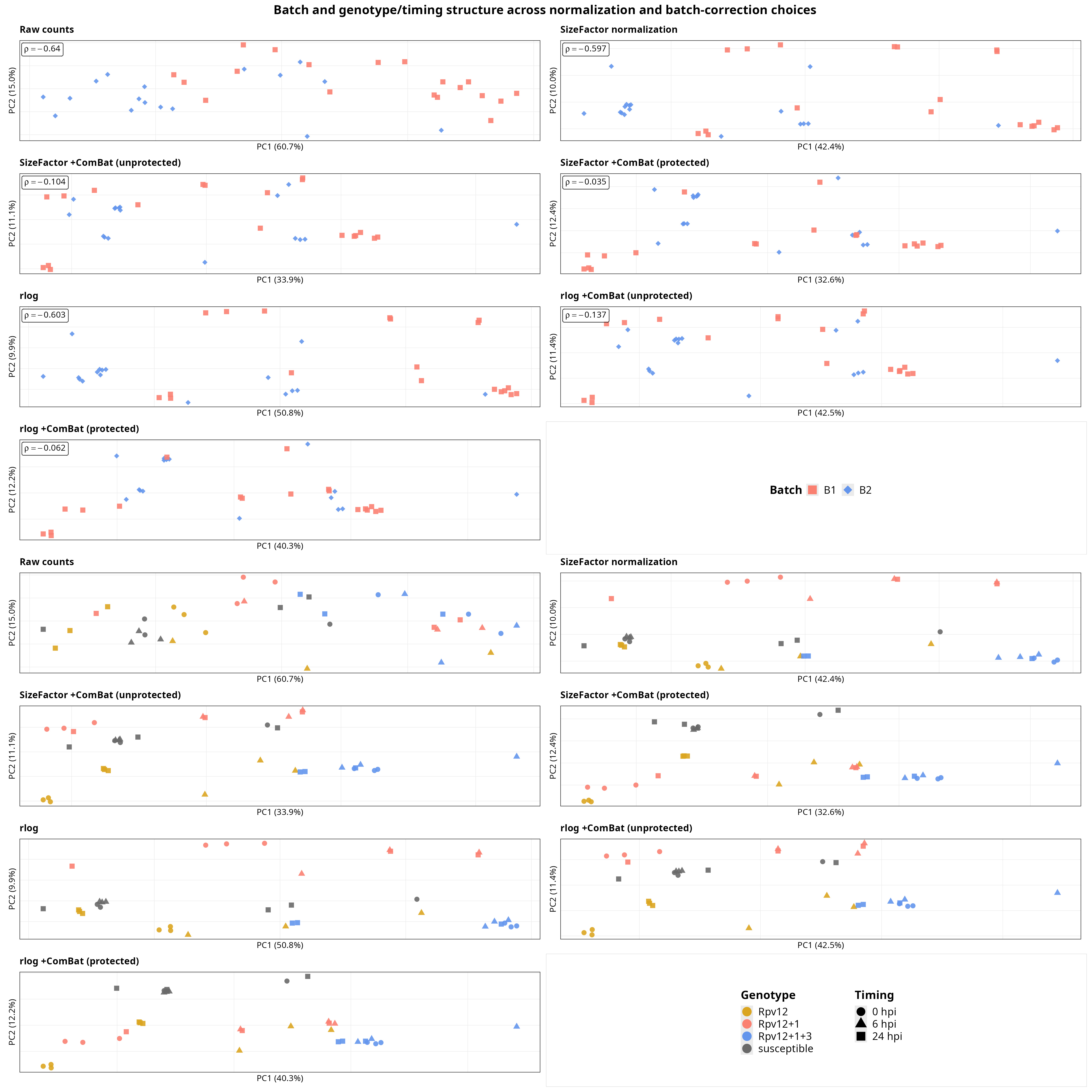

### Supplementary Figure 3

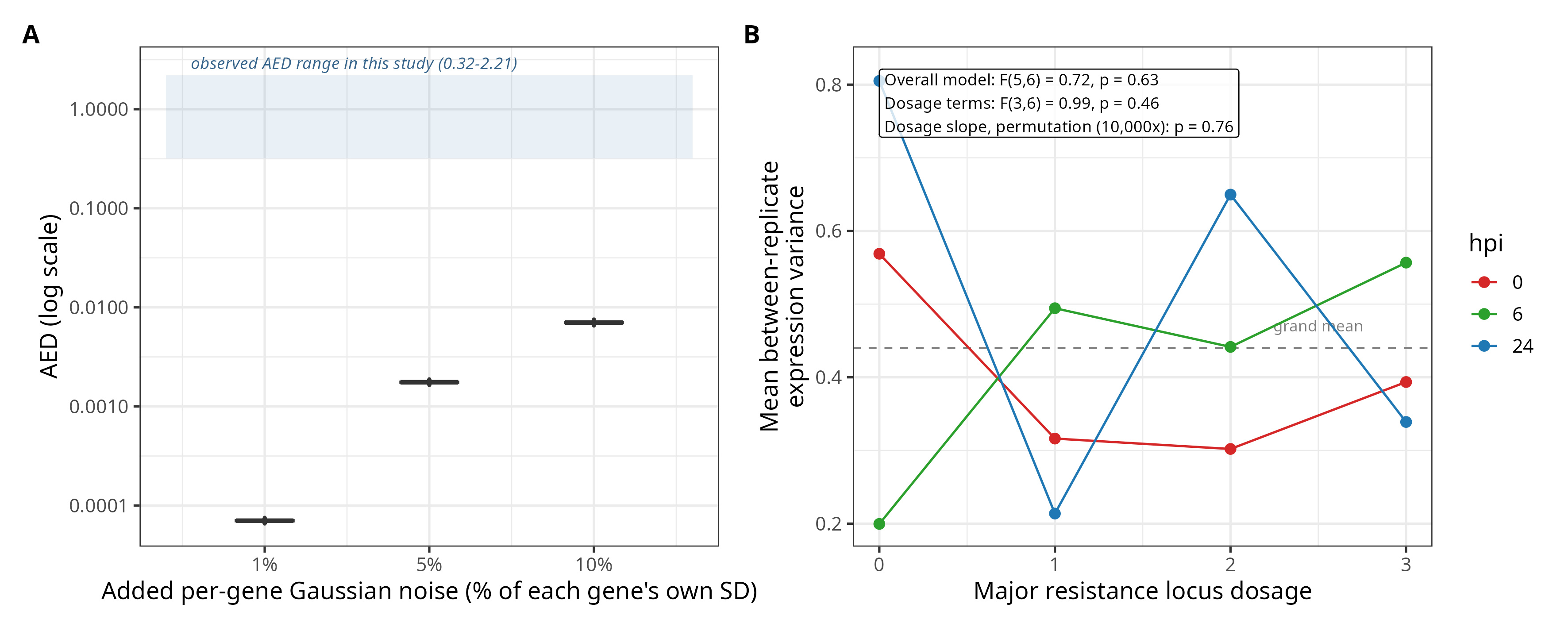

### Supplementary Figure 4

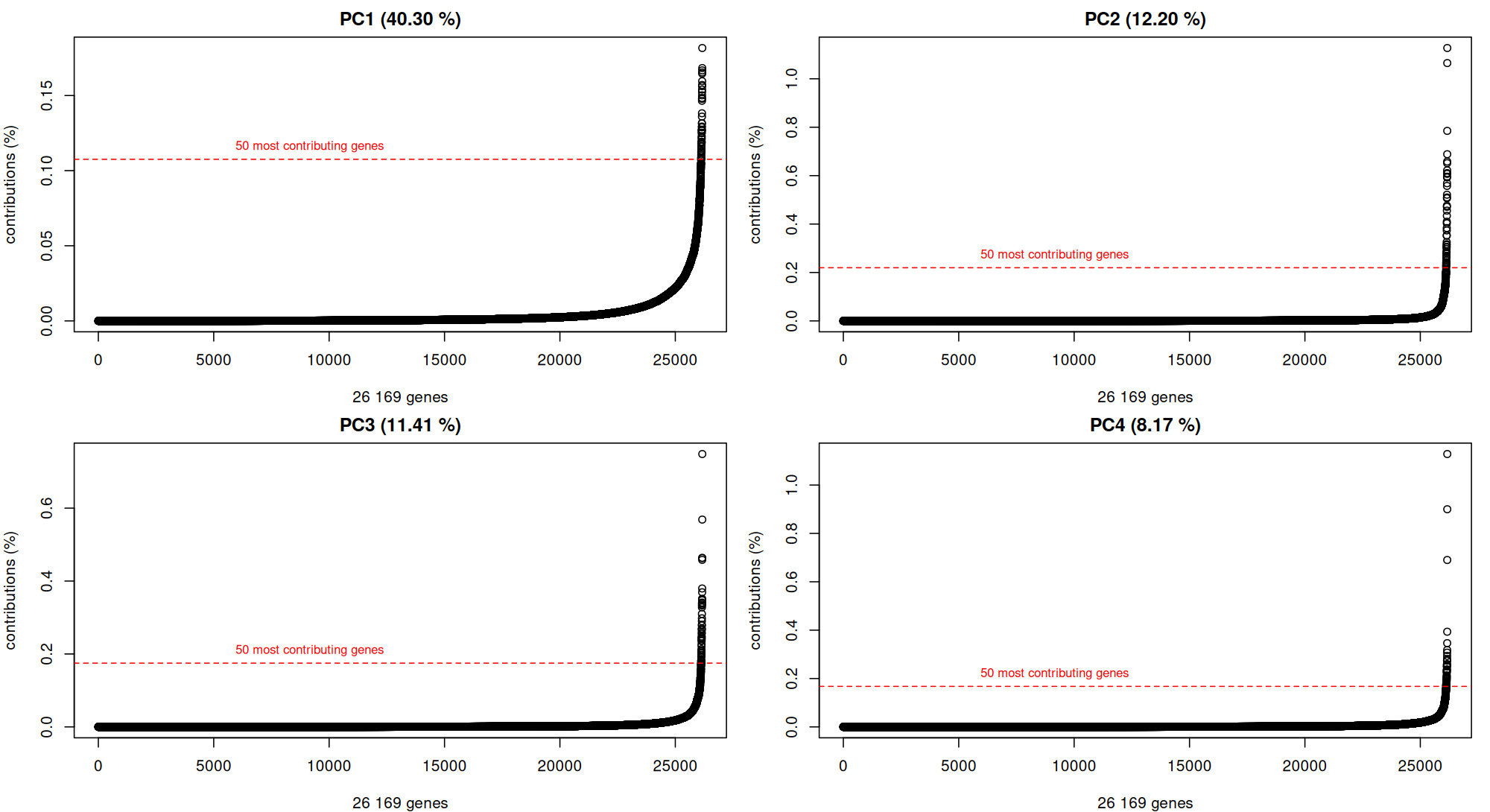

### Supplementary Figure 5

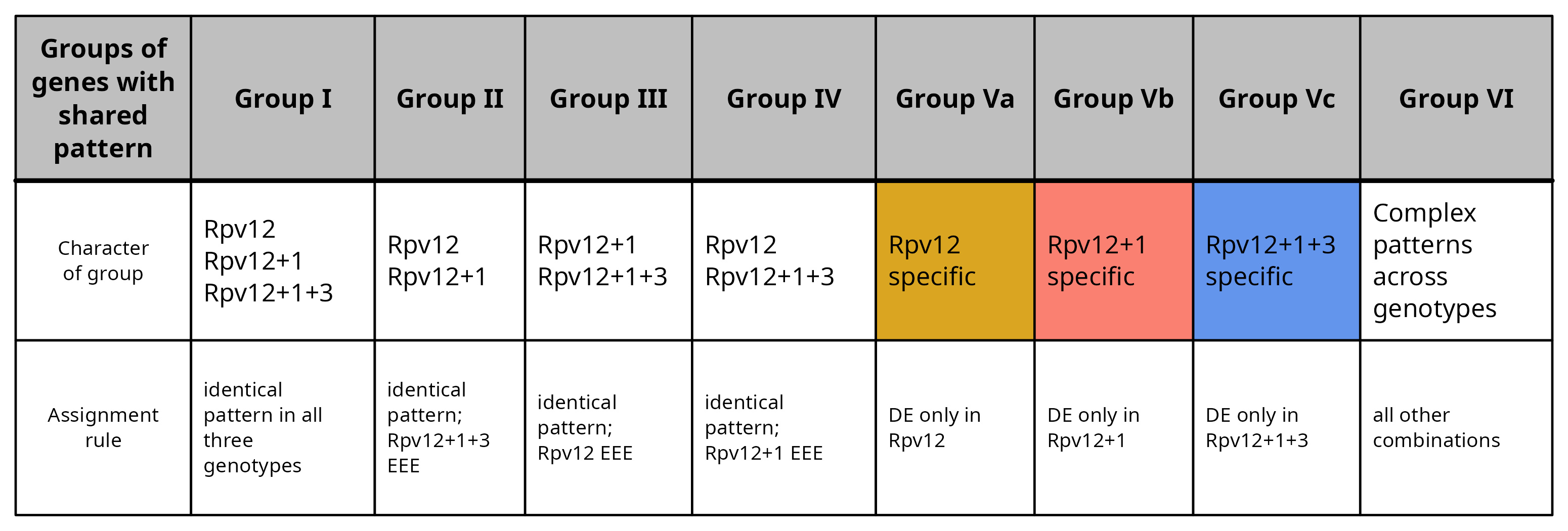

### Supplementary Figure 6

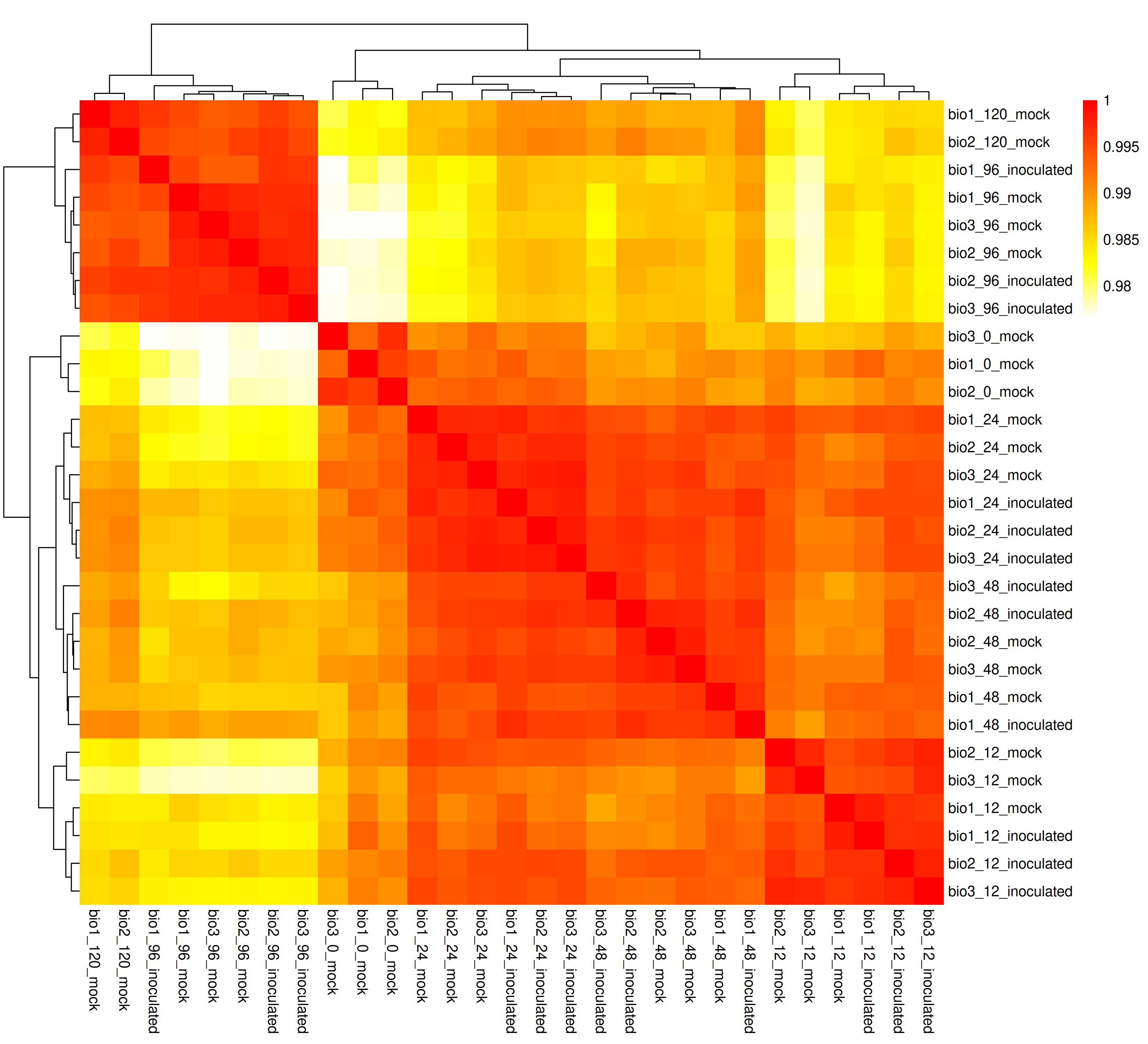

### Supplementary Figure 7

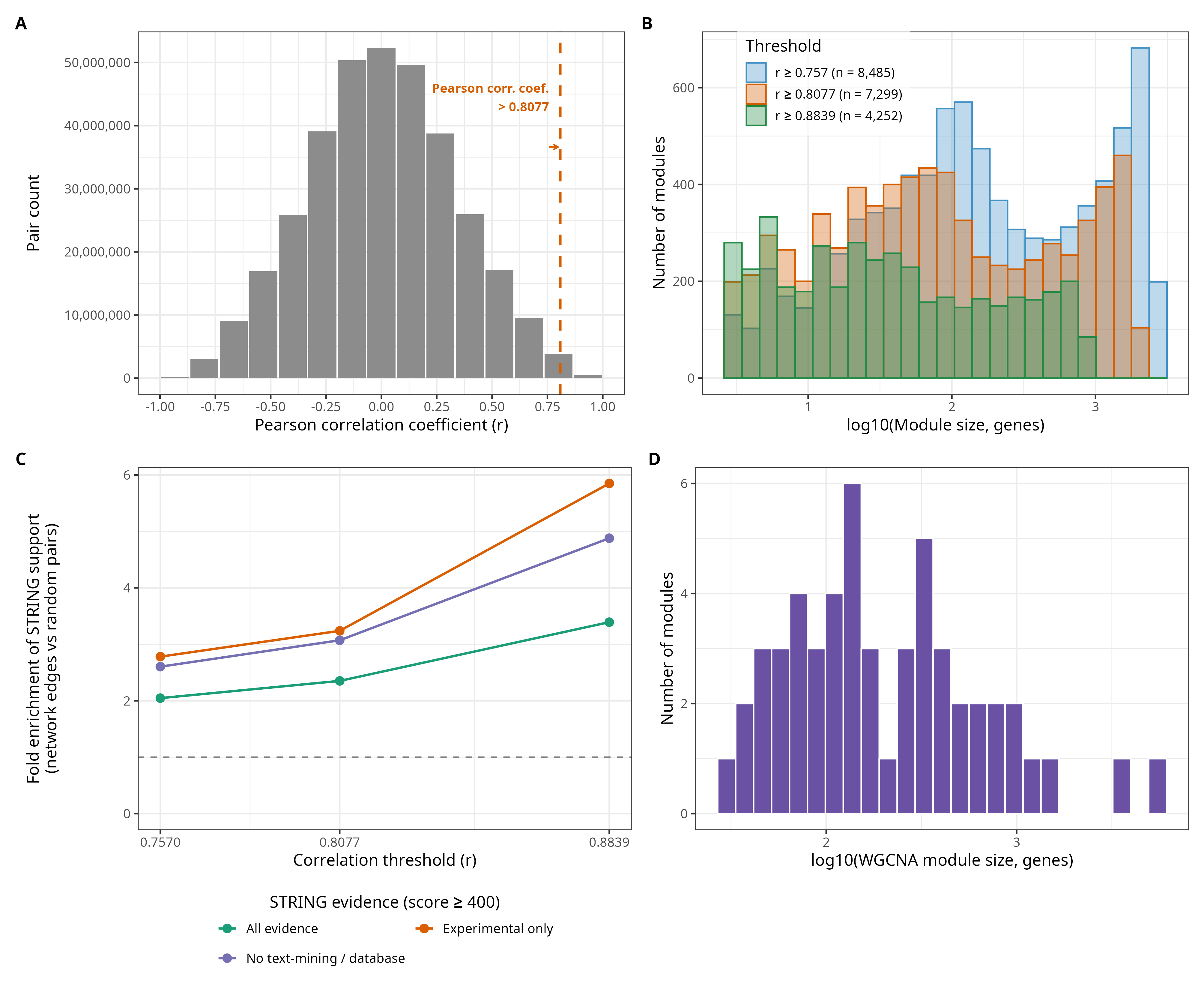

### Supplementary Figure 8

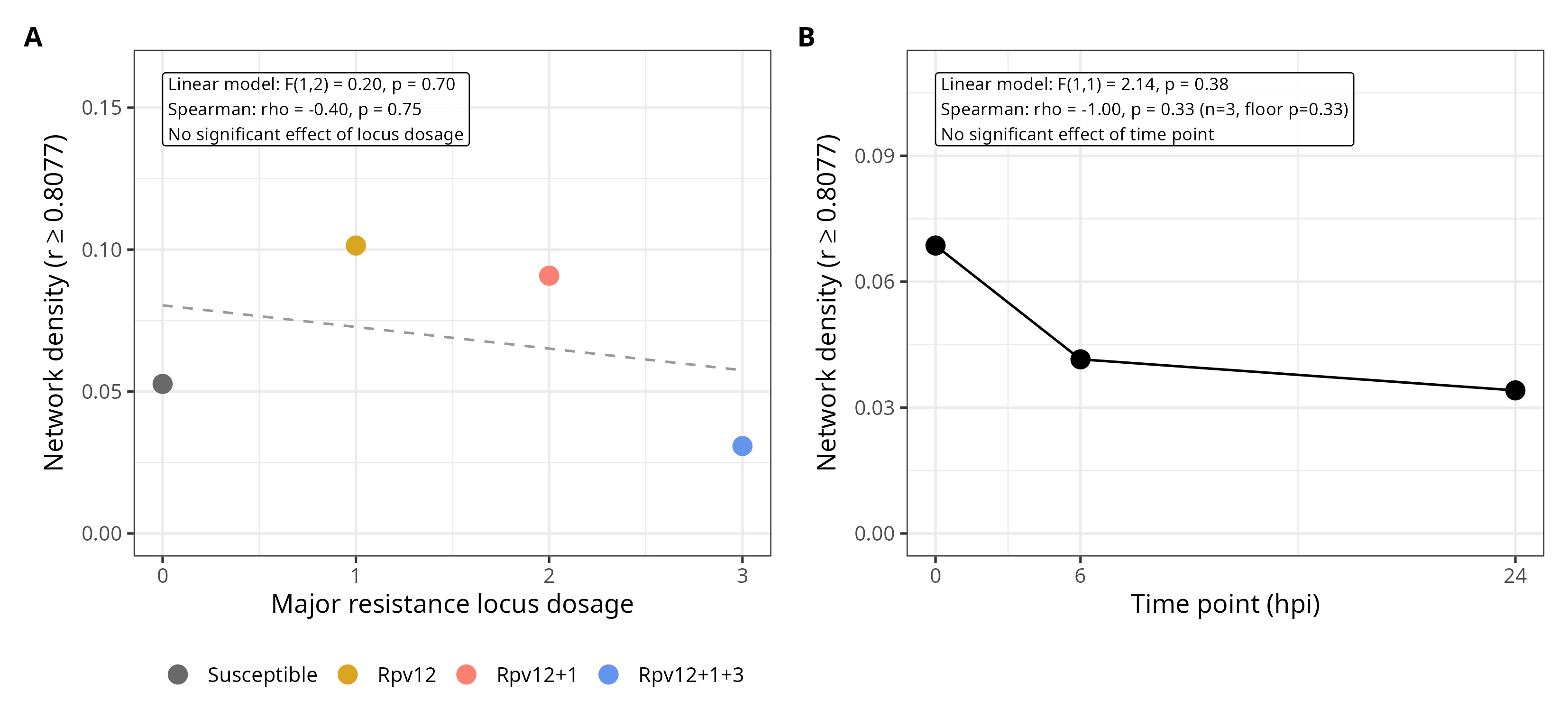
